# *Agropyron cristatum* genome provides a new insight for wheat improvement with wild relatives

**DOI:** 10.1101/2025.06.24.660289

**Authors:** Shenghui Zhou, Wenjing Yang, Haiming Han, Peng Zhou, Baojin Guo, Jinpeng Zhang, Bing Han, Xuezhong Liang, Yida Lin, Yuan Deng, Bin Liu, Xinming Yang, Yuqing Lu, Jisen Zhang, Ping Yang, Keke Yi, Yunbi Xu, Zhiyong Liu, Nils Stein, Qiang He, Hong-Qing Ling, Lihui Li

**Author notes:** These authors contributed equally (Shenghui Zhou, Wenjing Yang). Correspondence to Lihui Li, Hong-Qing Ling, Qiang He or Shenghui Zhou.

## Abstract

Exploring novel genetic variation for target traits is crucial for advancing wheat breeding and genetic improvement. Since 1990, we have been transferring genes from *Agropyron cristatum,* a promising wild relative genetic resource featuring robust spike morphology and stress tolerance, into common wheat by distant hybridization. To efficiently harness *A. cristatum* genes, we *de novo* assembled a 25.62-Gb high-quality genome of an autotetraploid accession Z559 with 28 chromosomes, and uncovered its specific genome features and an extensive repertoire of genes associated with yield and resistance to abiotic and biotic stresses. We systematically characterized *A. cristatum* genome introgressions by resequencing 431 wheat–*A. cristatum* derivatives and cloned the gene *AcGNS1*, which functions in the regulation of grain number per spike in wheat. Subsequently, we developed an optimized genome-selection model for wheat–*A. cristatum* breeding derivatives, enabling the rapid selection of new potential varieties. These findings demonstrate a genomic breeding strategy for leveraging novel genetic resources of wild relatives and advancing wheat improvement.

## Introduction

Common wheat (*Triticum aestivum* L.) is one of the agricultural founder crops domesticated during the Neolithic in the Fertile Crescent. Today, it is the most widely cultivated crop in the world, providing 20% of the human caloric intake (http://www.fao.org/faostat/en/, accessed 2024). Constant improvements in its grain yield and resistance to biotic and abiotic stresses are necessary for accommodating the needs of an expanding population and increasing food consumption as well as for adaptability of the climatic changes^1^. However, the narrow genetic diversity of wheat strongly limits the ability to breed varieties with significantly enhanced yield potential or adaptability to future climatic and agronomic challenges. The use of wheat wild relatives has played a crucial role in breeding novel wheat varieties^2^. Further discovery and transfer of favorable genes from wild relatives is crucial for broadening the genetic base and accelerating wheat breeding and genetic improvement.

*Agropyron* Gaertn. is a genus in the tribe Triticeae containing the P genome^3^. *A. cristatum* is a model species of this genus and widely cultivated as a high-quality pasture grass in many countries of the world^4^. More importantly, its exceptional spike morphology^5^—featuring 30 to 48 spikelets per spike and 6 to 12 florets per spikelet—along with desirable plant architecture and strong resistance to biotic and abiotic stresses^6^, positions it as a promising donor for wheat breeding in inter-species hybridization. Efforts to introgress traits from *Agropyron* into wheat began in the 1930s, but early attempts were unsuccessful^7,8^. A breakthrough was achieved in 1990 with the hybridization between common wheat cv. Fukuhokomugi (Fukuho) and a tetraploid *A. cristatum* accession Z559 (2n=4x=28, PPPP) from the Gobi Desert in Xinjiang, China^9^. This achievement initiated a new phase for extensive research on molecular cytogenetics and phenotypic analysis, resulting in the development of diverse wheat-*A. cristatum* derivatives, including addition, translocation^10,11^, and cytologically undetectable cryptic introgression lines^12–15^. These derivatives exhibited favorable agronomic traits, underscoring their strong potential for wheat improvement.

To address effective utilization of the genes of *A. cristatum* for wheat variety development, we assembled a high-quality genome of the tetraploid *A. cristatum* accession Z559 with PacBio HiFi reads, offering insights into its key genomic features and gene repertoire of its agronomic important traits. Through genetic analysis of 134 wheat-*A. cristatum* addition and translocation lines with the *A. cristatum* reference genomes, we dissected the P genome components introgressed into each line and rapidly identified and cloned a key gene *AcGNS1* from this wild species, which functions in the control of grain number per spikelet in wheat. Leveraging the information of introgressed elements and SNP variations of the 297 breeding derivatives, we developed a predictive breeding selection model, which is able to accelerate the selection of candidate varieties with high and stable yield. This study highlights the transformative potential of *A. cristatum* introgressions in wheat breeding.

## Results

### Genome assembly, validation and annotation

The genome size of *A. cristatum* accession Z559 was estimated to be approximately 24.40 Gb, nearly four times of 1C genome size of diploid *A. cristatum* (6,352 Mb)^16^, with a heterozygosity rate of 1.85% based on *k*-mer analysis (**Fig. 1a, Extended Data Fig. 1a and Supplementary Table 1**). Given its high heterozygosity and complex and tetraploid genome, we generated ∼624.41 Gb (30×) of PacBio HiFi reads with an N50 of 16.29 kb and 1,537 Gb (60×) Hi-C reads (**Supplementary Tables 2 and 3**). Initially, we assembled the Z559 genome using Hifiasm^17^, resulting in 25.62 Gb contigs with a contig N50 of 26.22 Mb (**Extended Data Table 1**). We then scaffolded the tetraploid genome by integrating 5.1 billion pair-reads of Hi-C data using ALLHiC^18^ and successfully anchored 24.37 Gb (95.12%) of the contigs to 28 pseudochromosomes, comprising seven homologous groups (HGs) with four haplotypic chromosomes per group (**Fig. 1b, Extended Data Table 1, Extended Data Fig. 1b and Supplementary Table 4**).

**Fig. 1:**
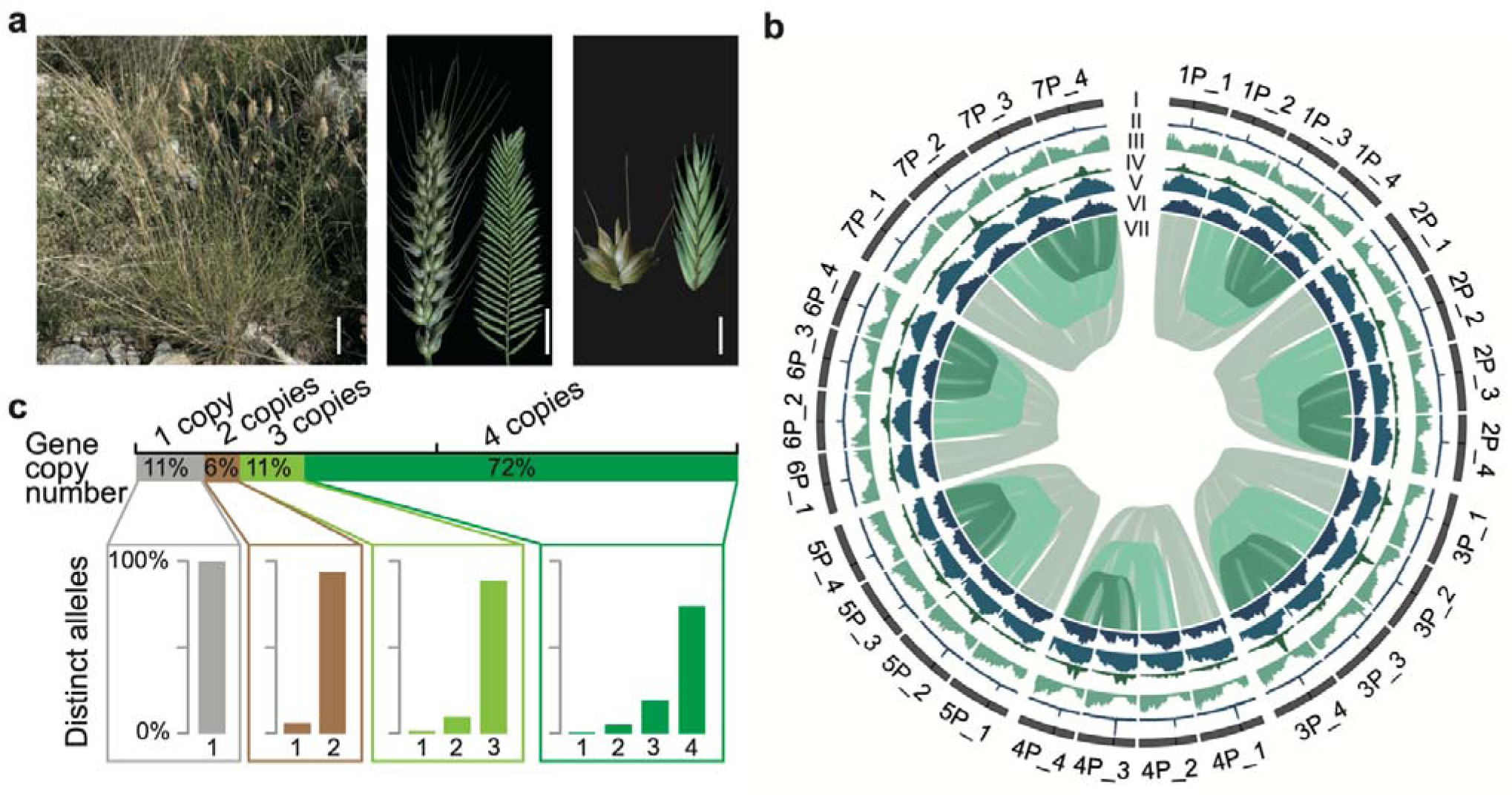
Overview of morphological characterization and genome of *A. cristatum* Z559. **a,** Morphological characterization of *A. cristatum* Z559, which was used for genome sequencing, in comparison with common wheat. Left, a representative niche of *A. cristatum*—the arid Gobi Desert (Zhou et al., 2017). Middle, spike morphology. Right, spikelet morphology. In the images of the spike and spikelet, the left side represents common wheat, while the right side represents *A. cristatum* Z559. Scale bars, 10 cm (left); 3 cm (middle); 1 cm (right). **b,** Circular representation of genomic features of *A. cristatum* Z559. The tracks show: I, chromosomes with functional centromere represented by black bar. II, CENH3 ChIP–seq coverage; III, gene density. IV, expression breadth density. V, density of *Gypsy* retrotransposons. VI, density of *Copia* retrotransposons. VII, syntenic gene pairs identified between chrX_4 and chrX_1, chrX_2, chrX_3. **c,** Allelic presence/absence variation of genes in the tetraploid genome of *A. cristatum* Z559. The gene copy number represents the number of alleles that each allele has. Distinct alleles refer to the number of distinct coding sequences of alleles with different copies.

To evaluate the completeness and accuracy of the Z559 genome assembly, we first mapped 1.10 Tb of Illumina short reads to the assembled genome (**Supplementary Table 5**). The mapping ratio reached 99.31% with a nucleotide accuracy approaching 100% (**Supplementary Table 6**) and a *k*-mer-based consensus quality value (QV)^19^ of 41.81 (**Table 1**), indicating that assembly accuracy of the Z559 genome is high. Additionally, we used the Benchmarking Universal Single-Copy Orthologs (BUSCO) method^20^ to evaluate gene completeness. The results showed that the BUSCO completeness of its four pseudo-monoploid genomes, each containing seven chromosomes, was 97.70%, 97.40%, 97.30%, and 96.50%, respectively, with a combined score of 98.70% (**Extended Data Table 1 and Supplementary Table 7**). The Long Terminal Repeat (LTR) Assembly Index (LAI)^21^ value reached to 22.96, which is the highest LAI reported among Triticeae genomes^22–34^ (**Extended Data Fig. 1c and 1d and Supplementary Table 8**). Chromatin immunoprecipitation with sequencing (ChIP-seq) analysis of centromere-specific histone H3-like protein (CENH3)^35^ further confirmed that the functional centromeric regions were contiguously assembled without sequence gaps (**Fig. 1b and Extended Data Fig. 2**). All these results demonstrate that the assembly of *A. cristatum* Z559 genome is of high quality (**Extended Data Fig. 1d**).

By integrating RNA (**Supplementary Tables 9**) and full-length transcriptome sequencing read alignments with *ab initio* and protein homology-based predictions (**Supplementary Tables 10**), we annotated 171,519 high-confidence (HC) protein-coding gene models across the four haplotypes of the *A. cristatum* Z559 genome (**Extended Data Table 1**). The overall BUSCO completeness score of 97.5% (1,574 out of 1,614 embryophyta single-copy orthologs) (**Extended Data Table 1 and Supplementary Table 7**) supports the predicted gene models with a high quality. In addition, we annotated 86.81% of the genome as transposable elements (TEs). Of them, LTR retrotransposons (LTR-RTs) are mostly abundant, accounting for ∼76.37% of the genome (**Supplementary Table 11**). Among these, the *Gypsy* and *Copia* are two major families, occupying 50.18% and 18.23% of the genome, respectively. Similar as Triticeae genomes reported previously, the genes, TEs, transcripts and expression breadth of the *A. cristatum* Z559 genome are not evenly distributed along the chromosomes^34^. The gene and transcript density were increasing on both arms along the centromere-telomere axis, but the TE and expression breadth showed the opposite pattern, correlating with the distance to centromere (**Fig. 1b**)^34^.

### Characteristics of the P genomes in the tetraploid *A. cristatum*

Among the HC gene models, 40,445 allelic genes were well-defined, 29,129 (72.02%) of which were present in all four haplotype genomes (quadruplex) (**Fig. 1c and Supplementary Table 12**), whereas the remaining alleles were present in three (triplex, 4,423, 10.94%), two (duplex, 2,298, 5.68%), or one haplotype genome (singleton, 4,595, 11.36%), indicating abundant allelic presence/absence variation across the four haplotypic genomes. Additionally, 3,194 tandem (consecutive) and 18,151 dispersed duplications were identified using allele-defined gene models, revealing extensive allelic copy number variation. On average, each gene has 3.44 copies and 3.18 distinct alleles in the accession Z559 genome (**Fig. 1c and Supplementary Table 12**).

To investigate the genomic divergence within the polyploid genome of *A*. *cristatum*, we first compared the coding sequences (CDS) of allelic gene pairs across the four haplotypic chromosomes. The analysis revealed a high CDS similarity (99.4%) with no significant differences between chromosomal counterparts (Student’s *t*-test *P* value > 0.05; **Extended Data Fig. 3a**). Phylogenetic analysis indicated that the four homologous chromosomes of *A. cristatum* shared a common origin (**Extended Data Fig. 3b**). Using the wheat D subgenome and the barley genome as references, we further assessed the *Ka*:*Ks* ratios for orthologous genes, and find no significant differences between chromosomal counterparts (Student’s *t*-test *P* value > 0.05; **Extended Data Fig. 3c and Supplementary Fig. 1**). These results suggest a symmetric purifying selection across all four haplotype genomes of *A. cristatum*. Collinearity analysis also revealed minimal structural divergence among the chromosomes (**Fig. 1b and Supplementary Fig. 2**). Gene expression analysis displayed similar overall expression levels across the homologous chromosomes (**Extended Data Fig. 3d**), indicating no significant genome dominance. Comparative analysis of syntenic gene pairs between monoploid genomes of autopolyploids and allopolyploids (**Supplementary Fig. 3**) placed *A. cristatum* clearly within the autopolyploid group, exhibiting the lowest average *Ks* value and the highest sequence identity (∼99%) among all autopolyploids analyzed (**Extended Data Fig. 3e**). These findings provide robust genomic evidence supporting a single origin of the P genomes in *A. cristatum*, resolving the longstanding debate on whether it is an auto- or allopolyploid, as previously speculated based on cytogenetic data^36^.

### P genome evolution

To elucidate the evolutionary trajectory of the *A. cristatum* P genome within the Triticeae, we conducted phylogenetic analysis using 1,928 single-copy orthologs across 14 genomes/subgenomes across the grass family. Our results indicate that the divergence between *A. cristatum* and diploid wheat occurred after the separation of barley but before the divergence of rye and *Thinopyrum elongatum* from wheat (**Fig. 2a**). The *Ks* distribution of the orthologous gene pairs suggests that the divergence of *A. cristatum* and wheat from their most recent common ancestor (MRCA) occurred approximately 8.80∼9.02 million years ago (Mya), and that tetraploidization of *A. cristatum* took place 0.95 Mya (**Extended Data Fig. 4a**).

**Fig. 2:**
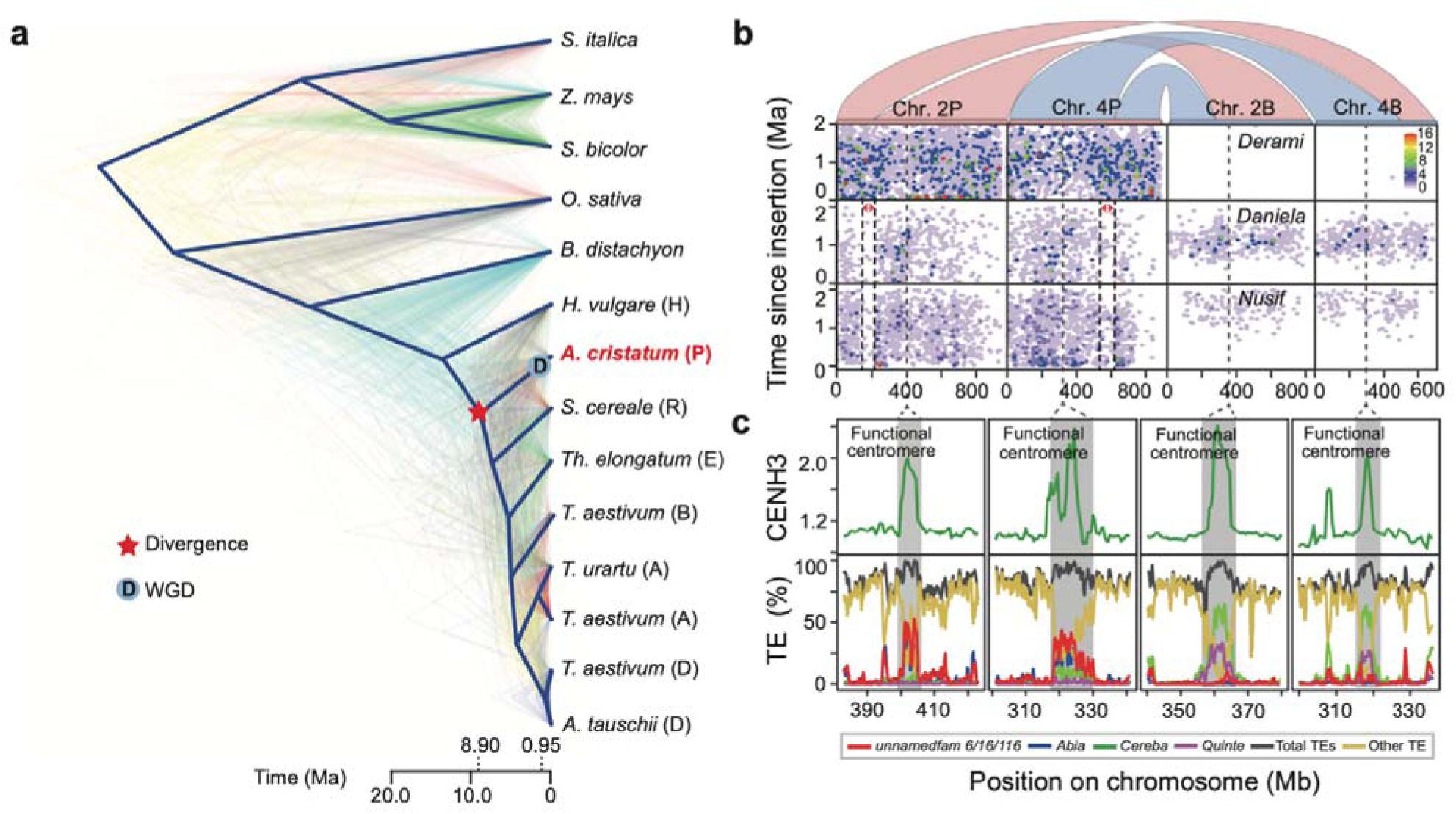
Genomic features of *A. cristatum* Z559 genome. **a,** Phylogenetic analysis of *A. cristatum* Z559 genome based on a gene tree constructed from 1,928 single-copy orthologous genes shared between the *A. cristatum* genome and 13 other grass genomes/subgenomes. The asterisk indicates the divergence of *A. cristatum* and the corresponding time, while “D” denotes the polyploidization of *A. cristatum* and its corresponding time. **b,** Syntenic blocks and distribution of representative three full-length LTR-RT families, including their positions and ages, between chromosomes 2P_1 and 4P_1 of *A. cristatum* Z559 and chromosomes 2B and 4B of the *T. aestivum* B genome. A representative inversion fragment is marked in blue, while chromosomal translocations are highlighted in red. The vertical axis represents binned ages (from 0 million years ago [Ma] at the bottom), and the horizontal axis represents bin positions. The heatmap indicates the number of LTR-RTs in each age/position bin (see legend inset). Red arrows highlight notable changes in LTR-RT profiles. **c,** Centromere composition of *A. cristatum* chromosomes 2P_1 and 4P_1 compared with *T. aestivum* chromosomes 2B and 4B. The top track shows CENH3 ChIP–seq coverage, while the bottom track depicts TE composition. Chromosomal positions are shown on the x-axis in megabases (Mb). Functional centromeres are indicated with shading.

Chromosomal rearrangement plays an important role in eukaryotic speciation at a macroevolutionary level^37^. Gene collinearity plots between *A. cristatum*, rye^31^, barley^28^ and wheat^34^ exhibited extensive genome collinearity (**Extended Data Fig. 4b-d**). However, we also found a large number of structural rearrangements in different chromosomes compared with the wheat, rye and barley chromosomes. The formation of 4P and 2P chromosomes, for example, has undergone chromosome fission, translocation and fusion (**Fig. 2b and Extended Data Fig. 4b-d**).

### TE composition of the *A. cristatum* genome

LTR retrotransposons (LTR-RTs) represent the largest proportion of most plant genomes and play significant roles in shaping genome structure and functions^38^. In the *A. cristatum* genome, 24 full-length LTR-RT families, such as *Deramin* and *Nusif*, have undergone substantial expansion (**Fig. 2b and Supplementary Table 13**). These LTR-RT families have undergone continuous expansion over the past ∼2 million years in the *A. cristatum* genome, whereas in the compared genomes, such as barley, rye, *Th. elongatum*, and wheat, they are either absent or show only episodic and limited activity (**Fig. 2b and Extended Data Fig. 5**). The sustained amplification of these LTR-RTs explains the prolonged retrotransposon burst observed in the *A. cristatum* genome (**Supplementary Fig. 4**).

Certain LTR-RT families, such as *Daniela* and *Nusif*, exhibit consistently lower abundance in peri-telomeric regions compared to other chromosomal regions (**Fig. 2b and Extended Data Fig. 5**). Interestingly, a similar scarcity was observed in the fusion regions of structural rearrangements on chromosomes 2P and 4P (**Fig. 2b**). These results suggest that the burst of *Daniela* and *Nusif* in *A. cristatum* genome exhibits a conserved expansion preference because the fusion regions on 2P and 4P were originally in the peri-telomeric regions before their rearrangement (**Fig. 2b and Extended Data Fig. 4b-d**).

LTR-RTs are major components of centromeres, essential for maintaining chromosomal integrity during cell division^39^. The functional centromere is defined by the epigenetic replacement of histone H3 with CENH3^35^. Through ChIP-seq analysis of CENH3, we identified distinct high-coverage region on each chromosome corresponding to the functional centromere in the *A. cristatum* Z559 genome (**Extended Data Fig. 6**). The centromere length ranged from 5.5 Mb on chromosome 1P_4 to 10.5 Mb on chromosome 5P_4 with 7.5 Mb on average (**Supplementary Table 14**). Unlike wheat and einkorn, where the *Gypsy* family elements *Cereba* (approximately 70%) and *Quinta* (approximately 20%) dominate centromeric regions, a unique composition pattern with approximately 40% *unnamedfam6* and 20% *Abia* elements was observed in the centromeric regions of *A. cristatum* genome (**Fig. 2c and Extended Data Fig. 6**). Moreover, we amplified an 871 bp sequence partially homologous to the *unnamedfam6* element in *A. cristatum*, which allowed clear differentiation of the P genome from other Triticeae genomes through fluorescent *in situ* hybridization (FISH) (**Supplementary Fig. 5**). These findings highlight unique features of the P genome centromeres, offering valuable insights into the evolutionary mechanisms underlying centromere function.

### Expansion of agronomically important gene families in *A. cristatum* genome

Genes undergoing significant alterations can play a large role in speciation, and may partly account for its phenotypic differentiation^40^. We performed comprehensive comparison on gene families among *A. cristatum*, rye, *Th. elongatum*^25^, and wheat, and identified 2,335 *A. cristatum*-specific gene families (comprising 8,025 genes). Functional enrichment analysis revealed that these specific genes were significantly enriched in essential Gene Ontology (GO) terms such as ion channel activity, ion transport, and enzyme inhibitor activity (**Supplementary Fig. 6 and Supplementary Table 15**), as well as in Kyoto Encyclopedia of Genes and Genomes (KEGG) pathways related to plant–pathogen interactions, photosynthesis, and the biosynthesis of secondary metabolites (**Supplementary Fig. 6 and Supplementary Table 16**).

Subsequently, we identified 2,511 rapidly expanded gene families (comprising 56,383 genes) in *A. cristatum*. Trait Ontology (TO) analysis of these rapidly expanded genes suggest that they are mainly associated with important agronomic traits, including resistance, rooting depth, grain number per plant and spikelet number per spike (**Supplementary Fig. 7 and Supplementary Table 17**). For example, *SPL14,* involved in rice yield and immunity to rice blast^41,42^, has only one copy in the A, B, D subgenome of common wheat, R genome of rye and E genome of *Th. elongatum*, but there are 16, 7, 3 and 10 copies in the four haplotypes of chromosome 7P, respectively (**Extended Data Fig. 7a and Supplementary Table 18**). The gene *DREB1C*, whose overexpression in rice resulted in 41.3 to 68.3% higher yield than wild-type due to increased grain number per panicle, grain weight, and harvest index^43^, expanded for 4 to 8 copies in the four haplotypes of chromosome 7P, while it has only 1 to 4 copies in the A, B, D, R and E genomes, respectively.

Other key examples include the major *Fusarium* head blight (FHB) resistance gene *Fhb7* derived from *Th. elongatum* had no orthologous genes in the A, B, D and R genomes^25^, but five, two and one copies were detected on chromosome 7P_1, 7P_4 and 3P_1 of *A. cristatum*, respectively. In term of abiotic stress tolerance, *CBF* is a key component in cold signal perception and responsive pathway^44^ (**Extended Data Fig. 7b**). In addition to the *CBF* gene cluster at the *Fr2* homology locus on HG 5 of *A. cristatum*, we also found another P genome-specific cluster of *CBF*-related genes on 1P and 2P (**Extended Data Fig. 7c and supplementary Table 18**).

Furthermore, 696 genes under natural positive selection were identified in the *A. cristatum* Z559 genome, such as *COLD1* (**Extended Data Fig. 7d**), *LG1* and *SPY* (**Supplementary Table 19**). Notably, these positively selected genes were primarily enriched in TO terms related to key agronomic traits, including flowering time, grain yield per plant, root number, plant height, spikelet fertility, and grain number per spike (**Supplementary Fig. 8 and Supplementary Table 20**).

All the results suggest that the expansion of these genes may provide a genetic basis for many favorable traits observed in *A. cristatum* and help guide the use of this wild species in wheat improvement.

### Analysis of wheat-*A. cristatum* addition and translocation lines with the assembled *A. cristatum* genome

Alien addition and translocation lines serve as essential intermediate materials in introgression breeding, act as donors to broaden the genetic diversity of crops. Using the assembled *A. cristatum* genome, we delineated the introgressed genetic components from *A. cristatum* in 134 lines generated by the hybridizations between the common wheat variety Fukuho and *A. cristatum* accession Z559, including 46 whole-chromosome addition lines, 17 partial chromosome addition lines, and 71 translocation lines (**Fig. 3a, Extended Data Fig. 8 and Supplementary Table 21**). A total of 143 introgressed chromosome segments of *A. cristatum* were identified across these lines, encompassing 45,753 non-redundant genes, including 2125 *A. cristatum*-specific genes.

**Fig. 3:**
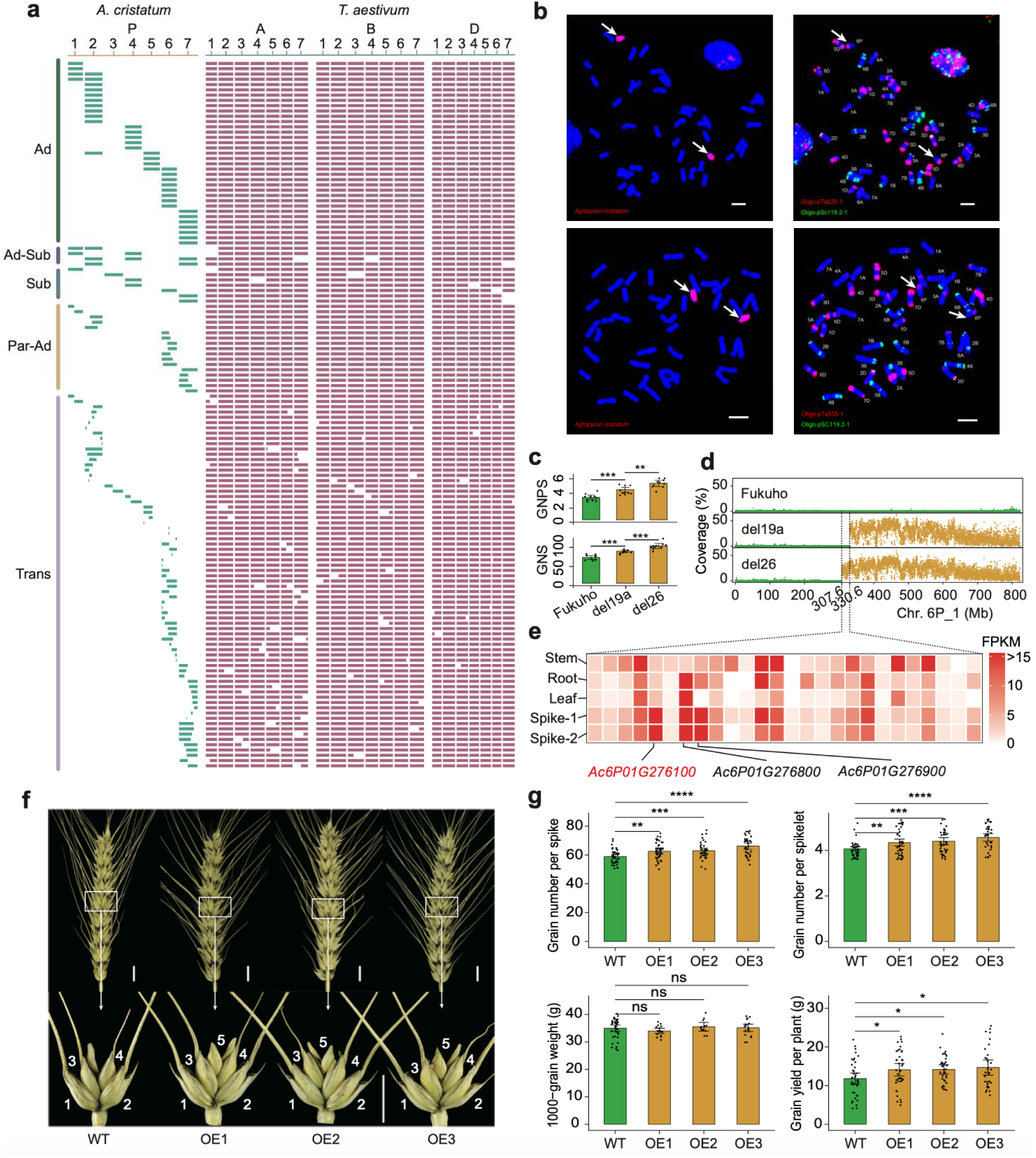
Genetic analysis of wheat-*A. cristatum* addition and translocation lines. **a,** Physical mapping of *A. cristatum* chromosomes/segments in 134 wheat-*A. cristatum* addition and translocation lines based on whole-genome sequencing. Each row represents a derivative line, with green blocks indicating *A. cristatum* genome chromatin and red blocks indicating wheat genome chromatin. Line categories: Ad (addition lines), Ad-Sub (addition and substitution lines), Sub (substitution lines), Par-Ad (partial chromosome addition lines), and Trans (translocation lines). **b,** Cytological identification of wheat-*A.* arrows indicate the chromosomes of *A. cristatum*. Scale bar: 10 μm. **c,** Statistical comparison of grain number per spikelet (GNPS) and grain number per spike (GNS) among Fukuho, del19a, and del26 lines. Error bars represent ± SE, and *P* values were calculated using a two-sided t-test. **d,** Resequencing coverage distribution along chromosome 6P_1 in wheat-*A. cristatum* partial addition lines del19a (middle) and del26 (bottom), compared to their wheat parent Fukuho (top). Coverage was calculated in 10 kb sliding windows. Windows corresponding to *A. cristatum* introgression blocks are highlighted in yellow. **e,** Heatmap of 26 genes expressed in the young spikes of addition line del26 within the 6PS_1 region (307.60–330.56 Mb). Young spike samples were collected at the double ridge stage (Spike-1) and spikelet differentiation stage (Spike-2). Numeric expression values for predicted genes are detailed in Supplementary Table 20. The candidate gene is highlighted in red. **f,** Phenotypic characterization of *AcGNS1*-overexpressing lines (OE1–OE3) compared to Fielder (wild type, WT). Top, spikes (scale bar = 1 cm). Bottom, spikelets (scale bar = 1 cm). **g,** Statistical comparison of yield-related traits between Fielder and the *AcGNS1*-overexpressing (OE) lines. Error bars indicate mean ± SE. Bars marked with * and ** denote significant differences between WT and the OE lines at *P* < 0.05 and *P* < 0.01, respectively, based on Student’s t-test.

As a representative example, the addition line 4844-12, containing 6P chromosome, exhibited significant increase of grain number per spike (GNS) and thousand grain weight (TGW), while enhancing resistance to multiple diseases^45^. Using this line, we developed seven partial chromosome addition lines and 26 translocation lines (**Supplementary Table 21**) for wheat improvement. With the genome assembly and annotation of *A. cristatum,* we detected that Chr6P_1 of *A. cristatum* was added in the line 4844-12. The added Chr6P_1 contains 5,541 HC genes along with 686 homologs of known functional genes. On this chromosome, a set of yield-related genes were identified, including 136 grain number-related homologs and 67 grain weight-related homologs, such as *RFT1*, *LRK1*, *D1*, *GW6a*, and *GW2*. Additionally, we identified 510 disease resistance-related homologs, including *Pi21*, *Pm21*, *Pm2*, *Xa21*, and *Lr13*, as well as 181 abiotic stress tolerance-related homologs on this chromosome.

Among the seven partial chromosome addition lines derived from 4844-12, two lines, del19a (2n = 44) and del26 (2n = 44) (**Fig. 3a and Supplementary Table 21**), exhibited significantly higher grain number per spike (GNS) and grain number per spikelet (GNPS), compared to the receptor variety Fukuho (**Fig. 3c**). Notably, del26 displayed a greater increase in GNS and GNPS than did del19a (**Fig. 3c**). Genomic analysis revealed that del19a carries a 492.01 Mb segment from 330.56–822.57 Mb of *A. cristatum* Chr6P_1, whereas del26 contains a slightly larger segment (514.97 Mb) from 307.60–822.57 Mb of Chr6P_1 (**Fig. 3d**). The added chromosomal difference between the two lines is a 22.96 Mb region (Chr6P_1:307.60–330.56 Mb), implying that this interval harbors gene(s) controlling GNS and GNPS. Within the 22.96 Mb region, 53 HC genes were annotated (**Supplementary Table 22**). For identification and cloning of the responsible gene, we performed RNA-seq analysis of del26 across tissues, including root, stem, leaf, and young spike, and found that 26 genes in the 22.96 Mb region expressed in young spike tissues, and only three of them displayed high expression during critical developmental stages at double ridge and spikelet differentiation (**Fig. 3e**). Among the three genes, only *Ac6P01G276100* has functional annotation, showing homology to *SWI3B* of *Arabidopsis thaliana*, a gene implicated in early embryonic development^46^. Therefore, we considered *Ac6P01G276100* as the candidate gene influencing GNS and GNPS and designated it as *AcGNS1*.

To validate the role of *AcGNS1* in regulating grain traits, we over-expressed it in Fielder, a cultivar of common wheat. Transgenic lines with *AcGNS1* expression (**Supplementary Fig. 9**) exhibited significant increase in GNPS (by 0.36 grains; P < 0.05) and GNS (by 5.02 grains; P < 0.01) in the T_3_ generation (**Fig. 3f and g**), resulting in an 18.09% elevation of grain yield per plant (P < 0.05; **Fig. 3g**). These results confirm that expression of *AcGNS1* from *A. cristatum* in wheat can indeed enhance GNPS and GNS, consequently improving grain yield.

Overall, our resequencing analysis of these addition and translocation lines developed through distant hybridization helps to elucidate the genetic composition of the P genome in these materials, and conclude that these materials are new genetic sources hosting a large number of favorable *A. cristatum* genes for high yielding, disease resistance and abiotic stress tolerance, which are required for developing future wheat.

### Analysis of wheat-*A. cristatum* breeding derivatives with the assembled *A. cristatum* genome

Since 1990, we have developed 297 breeding derivatives through intergeneric hybridization between wheat cv. Fukuho and *A. cristatum* Z559, followed by multi-generational backcross with Fukuho and other main cultivated varieties, combined with artificial selection for target traits (**Extended Data Fig. 8 and Supplementary Fig. 10**). During the 2021–2022 growing season, we conducted a phenotypic survey of 6 yield-related traits (yield per unit area, thousand-grain weight, spike number per unit area, grain number per spike, spikelet number per spike and total sterile spikelet number per spike) on 141 breeding derivatives, under plot planting conditions with actual production density in Xinxiang, Henan province (**Supplementary Table 23**). Their yields varied from 7,912.84 kg/ha to 11,917.19 kg/ha, with an average yield of 10,372.78 kg/ha. Among these 141 derivatives, 124 derivatives (87.94%) produced higher yields than the local dominant variety (Zhoumai 18, 9,517.25 kg/ha), and 66 derivatives (46.81%) exceeded it by more than 10% (**Supplementary Table 23**). As a result, these derivatives exhibited superior yield performance in the field, significantly outperforming the local control varieties.

We employed a *k*-mer-based approach to detect the genomic fragments introgressed from *A. cristatum* in all 297 breeding derivatives (see methods). Totally, 107,779 introgressed fragments have been identified from *A. cristatum* with a cumulative size of 13.88 Gb across all derivatives, and 996 of them are non-redundant introgression segments (**Fig. 4a and Supplementary Table 24**). The sizes of the single introgression segments range from 50.00 kb to 1.25LMb with 104.70 kb on average. Each derivative contained 131 to 500 introgression segments with an average number of 363 (**Fig. 4a and Supplementary Table 24**). The line with the largest total length of introgression segments is xq336, with a cumulative introgression size of 62.83 Mb. Interestingly, we found that the introgression segments are unevenly distributed across the wheat genome, and more introgression segments come from *A. cristatum* chromosome arms than that from centromeric regions (**Fig. 4a**). Out of the 996 non-redundant introgression segments, 351 (35.24%) contain protein-coding genes. Enrichment analysis revealed that introgressed genes are significantly enriched in GO terms such as protein glycosylation, enzyme inhibitor activity, KEGG pathways including cutin, suberine and wax biosynthesis, and TO terms related to grain yield per plant, grain number per plant and primary branch number, and others (**Supplementary Table 24 and Supplementary Fig. 11**).

**Figure 4:**
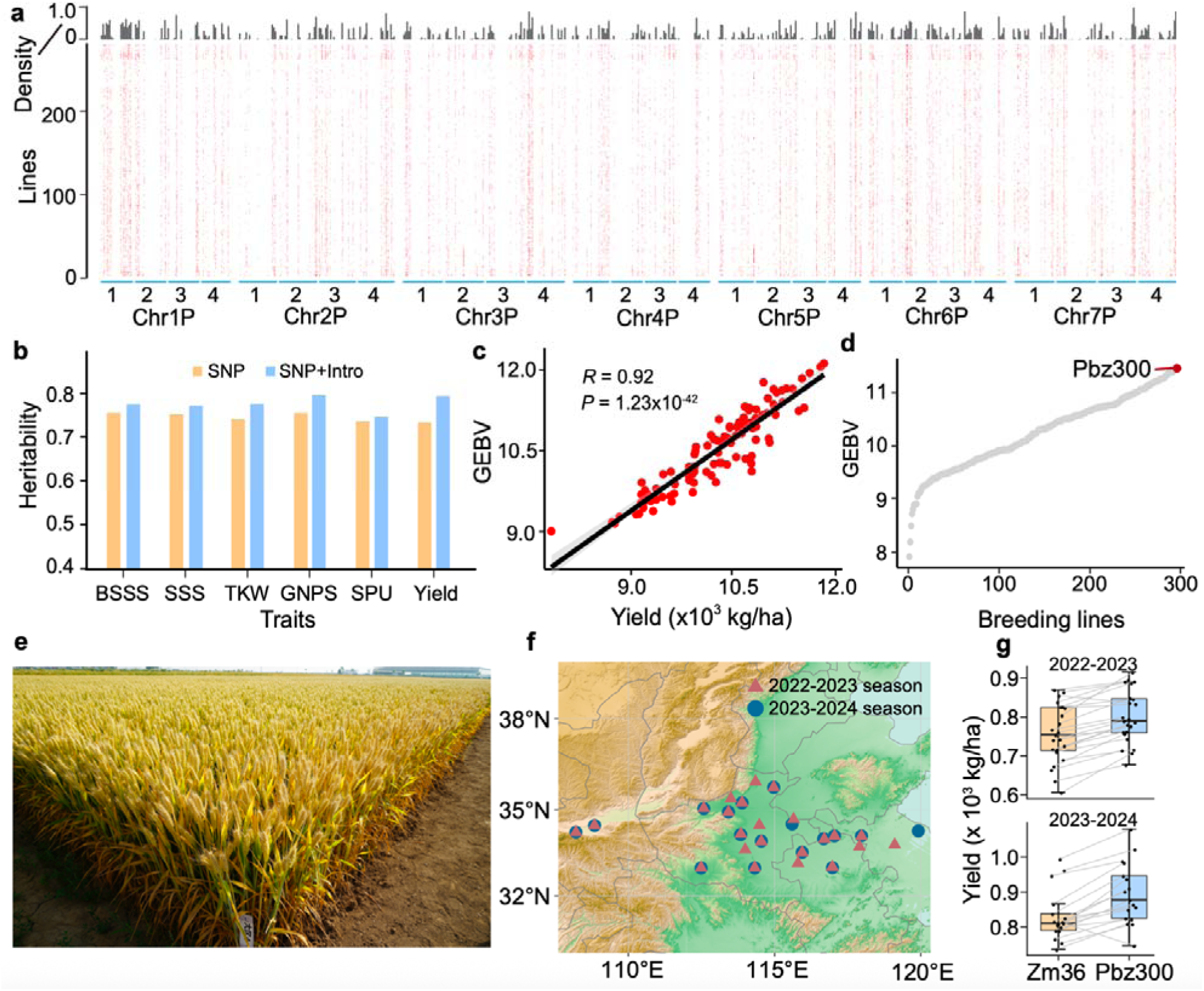
Genomic selection model developed in wheat-*A. cristatum* breeding derivatives accelerate the breeding of elite wheat varieties. **a,** Distribution of *A. cristatum* introgression segments across 297 wheat-*A. cristatum* breeding derivatives. The horizontal axis represents the 28 chromosomes of the *A. cristatum* Z559 genome, with 1-4 indicating four different haplotype chromosomes. On the vertical axis, ‘Lines’ represent the 297 wheat-*A. cristatum* breeding derivatives, while ‘Density’ indicates the proportion of breeding derivatives containing a specific introgression fragment relative to all breeding derivatives. **b,** Heritability estimates of six yield-related traits using wheat SNP genotypic data alone (“SNP”) or combined wheat SNP and *A. cristatum* introgression segment genotypic data (“SNP+Intro”). Traits include: SSNS (sterile spikelet number per spike), SNS (spikelet number per spike), TGW (thousand grain weight), GNS (grain number per spike), SNU (spike number per unit area), Yield (yield per unit area). **c,** Genomic selection model accurately predicted GEBVs of yield traits in the hold-out test dataset. **d,** Pubingzi 300 (Pbz300) showed the highest predicted GEBV among the 297 breeding derivatives. **e,** Field performance of Pbz300. **f,** National regional trial of Pbz300 conducted across 23 and 17 sites in the 2022-2023 and 2023-2024 seasons, respectively. **g,** ield

To evaluate the contribution of *A. cristatum* introgression to yield-related traits, we used a linkage-disequilibrium-adjusted kinships (LDAK) model (see methods) to analyze these phenotyped breeding derivatives, estimating the phenotypic variance explained by genetic variants, including *A. cristatum-*introgression polymorphisms and wheat single nucleotide polymorphism (SNP). The heritability value of the investigated 6 traits ranged from 0.71 to 0.74 without introgression information (i.e., SNPs only) (**Fig. 4b**). In contrast, incorporating introgression polymorphism information led to the significant increase in heritability for yield trait, with an improvement of 9.7%, and an average increase of 4.6% across the six traits (**Fig. 4b**).

We further developed genomic selection (GS) models using these phenotyped derivatives as the training population to predict the agronomic traits, aiming to rapidly select the candidate varieties or lines from the all 297 breeding derivatives for production. When only trait-associated SNP markers (see methods) were used, the average prediction accuracy for the 6 agronomic traits ranged from 0.558 to 0.822 (**Supplementary Fig. 12**). However, incorporating introgression segment markers into the models led to the greatest improvement in prediction accuracy for thousand grain weight, with an increase of up to 41.3%, and an average enhancement of 14.6% across all six evaluated traits (**Fig. 4c and Supplementary Fig. 12**), suggesting that the introgressed fragments from *A. cristatum* are positively involved in grain yield formation. To identify the derivatives with the greatest potential to become new varieties, we used the optimal yield GS model to predict the genomic estimated breeding values (GEBVs) for all 297 derivatives. As shown in **Fig. 4d**, the wheat-*A. cristatum* breeding derivative xq224, which contains 390 introgression segments, had the highest GEBV for yield among the 297 derivatives.

We then conducted national regional trials of the derivative xq224 (renamed as Pubingzi 300) at 23 sites in the Southern Huang-Huai region during the 2022-2023 growing season and at 17 sites during 2023-2024 growing seasons, according to the China New Wheat Variety Trial Specification (**Fig. 4e and f**). Pubingzi 300 exhibited outstanding performance with average yield of 8,005.5 kg/ha and 8,948.3 kg/ha in the two seasons. These yields were 5.86% and 7.84% higher, respectively, than the control variety Zhoumai 36 (China’s new variety approval standard mandates that yields must be at least 3% higher than those of the control variety). The highest yield improvement reached to 16.77% at the site Suzhou in 2022-2023, and 18.71% at the site Xuchang in 2023-2024. Notably, Pubingzi 300 exhibited stable performance across diverse environmental conditions, ranking first in average yield across all 17 test sites among the 19 candidate varieties during 2023-2024 (**Supplementary Table 26 and Fig. 4g**). These results highlight the potential of Pubingzi 300 as a highly promising wheat variety with high yield and broad regional adaptability, and further confirming the contribution of *A. cristatum* introgression to the enhancement of wheat yield.

## Discussion

Harnessing the genetic diversity of wild relatives remains a promising strategy to overcome the genetic limitations in modern crop breeding^2,47^. However, identification and utilization of agronomic important genes from wild relatives is an awe-inspiring task due to the challenges, such as low crossing achievement, high sterility, linkage drag, and suppressed recombination and a huge long breeding cycle with traditional distant hybridization. In this study, we successfully assembled a high-quality genome of *A. cristatum* (P genome), the wild donor species used in our distant hybridization program for wheat improvement^48–50^, and resequenced 431 wheat–*A. cristatum* derivatives. These genomic sequence resources combined with the predicted gene models provide a platform for precise identification of introgressed segments and genes from *A. cristatum,* offering a powerful framework to accelerate the utilization of *A. cristatum* genes for advancing wheat improvement with marker-associated selection. The sequence resource also opens the path towards identifying the underlying functional genes or regulatory sequences, which can be exploited for improvement of wheat and even other crops by gene transformation and genome editing approaches.

Comparative genome analyses revealed lineage-specific expansion of certain LTR-RT families, unique centromeric TE compositions, and genomic rearrangements, highlighting the evolutionary distinctiveness of the *A. cristatum* genome in Triticeae^23–32,34,51–53^. Importantly, the expansion of gene families related to spike development and stress responses suggests that *A. cristatum* harbors valuable agronomic potential beyond what is found in modern wheat.

By integrating this genome with resequencing data from wheat–*A. cristatum* derivatives, we revealed that these materials harbor not only functional genes underlying their own favorable agronomic traits but also a wide array of novel genes absent in wheat, which are associated with enhanced productivity, disease resistance, and abiotic stress tolerance. The stable integration of these alien-derived genes into wheat genome has substantially expanded the genetic base of modern wheat.

Notably, our assembly distinguishes the four haploid genomes of tetraploid *A. cristatum*, allowing haplotype-specific tracking of introgressions. However, its representativeness is inherently limited. Therefore, a *A. cristatum* pangenome will be essential for capturing broader haplotype diversity and unlocking the full potential of this species.

In summary, this work not only provides a foundational genomic resource of *A. cristatum,* but also introduces a rational breeding framework for exploiting wild germplasm. The ability to link introgressed segments to functional variation opens new avenues for trait improvement, including genome editing, and the strategies can be used in other crops.

## Methods

### Plant materials

The *A. cristatum* accession Z559 (2n = 4X = 28, PPPP), a tetraploid species native to the Gobi Desert in Xinjiang, China, was collected from one of the most expansive regions of *A. cristatum* distribution (AgroAtlas, 2015, http://www.agroatlas.ru/en/content/related/Agropyron_cristatum/). This accession is preserved in the National Crop Genebank of China. To introduce desirable traits from *A. cristatum* into wheat, wheat–*A. cristatum* Z559 alien whole chromosome addition lines were developed through hybridization between the common wheat Fukuho and *A. cristatum* accession Z559, followed by several generations of backcrossing or selfing. Partial chromosome addition and translocation lines were subsequently derived from these whole chromosome addition lines using ^60^Co-γ irradiation or by incorporating an *Aegilops cylindrica* gametocidal chromosome, followed by additional backcrossing to Fukuho (**Extended Data** Fig 8). Meanwhile, we developed a series of introgression lines that exhibited superior phenotypic traits specific to *A. cristatum*. To fully utilize the potential of these materials in breeding, they were subjected to multiple rounds of hybridization with other main wheat cultivars and selfing, followed by artificial selection for desirable agronomic traits since 1990. Ultimately, we successfully obtained 297 wheat–*A. cristatum* breeding derivatives (**Extended Data Fig 8 and Supplementary Fig. 10**).

### Illumina short reads sequencing

Genomic DNA was extracted from fresh leaf tissues of *A. cristatum* Z559 and all sequenced wheat and derivatives using the cetyl trimethyl ammonium bromide (CTAB) method^54^. 350-bp paired-end (PE) libraries were constructed using the NEBNext Ultra DNA Library Prep Kit. Sequencing was performed on the HiSeq 2500 platform (Illumina, San Diego, CA, USA) with a PE read length of 150 bp, and data generation was carried out by Novogene Technologies Co., Ltd. (Beijing, China). Raw sequencing data were processed using Trimmomatic^55^ to filter out low-quality reads and overrepresented sequences, ensuring high-quality downstream analysis.

### *A. cristatum* Z559 genome sequencing, assembly, and validation

#### Evaluation of genome size

Approximately 45× Illumina short reads were generated from three libraries to characterize the genome of *A. cristatum* Z559 and evaluate the genome assembly. The *k*-mer analysis was performed using Illumina DNA data prior to genome assembly to estimate the genome size and heterozygosity^56^. Briefly, the genome survey was performed through *K*-mer analysis using the software Jellyfish (v1.1.10)^57^ with parameter of ‘-m 19’ and GenomeScope (v2.0)^58^ with parameter of ‘kL=L19’.

#### Genome sequencing

Extracted DNA was used to construct circular consensus sequencing (CCS) libraries, which were sequenced on the PacBio Sequel platform. PacBio HiFi reads were generated using ccs software v.3.0.0 (https://github.com/pacificbiosciences/unanimity/, with parameters: --min-passes 3 --min-length 10000 --max-length 1000000 --min-rq 0.99) and used for the *de novo* assembly of the *A. cristatum* Z559 genome.

Fresh leaf tissues were fixed with 1% formaldehyde to prepare Hi-C libraries for sequencing. Following the protocol described previously^59^, chromatin was digested using HindIII and DpnII restriction enzymes. The libraries were then sequenced on the Illumina HiSeq platform, and the resulting Hi-C reads were used to construct and scaffold the genome assembly.

#### Genome assembly

To generate a high-quality genome assembly of *A. cristatum* Z559, we utilized PacBio HiFi sequencing data for primary assembly and chromosomal conformation capture (Hi-C) data for scaffolding. The haplotype-resolved *de novo* assembly was performed using hifiasm^17^, a specialized assembler designed for long, high-fidelity reads, with parameters -l 0 -k 51 -w 51 --write-paf --write-ec. Hifiasm first applied haplotype-aware error correction to retain heterozygous alleles while correcting sequence errors, and then constructed a phased assembly graph, connecting reads from the same haplotype and incorporating local phasing information. By integrating complementary data with global phasing information, hifiasm produced fully phased assemblies for each haplotype. To further scaffold the assembly, Hi-C reads were aligned to the contigs using HICUP (v0.7.3)^60^, and the ALLHiC algorithm^18^ was employed to cluster, order, and anchor the contigs into chromosomes. This process involved pruning interallelic links to separate homologous chromosomes, clustering contigs into groups based on Hi-C linkage signals, rescuing unpartitioned contigs using Hi-C signal density, optimizing the order and orientation of contigs, and finally building chromosomes by concatenating contigs and generating the final assembly in FASTA format. The genome was further refined using Juicebox Assembly Tools v1.11.08^61^ to visualize and manually correct large-scale structural variations, such as inversions and translocations, resulting in the final chromosome sequences.

#### Evaluation of the genome quality

The completeness of the *A. cristatum* genome assembly was evaluated using BUSCO against 1,614 embryophyte genes from the ‘Embryophyta_odb10’ dataset^62^. LTR_retriever^21^ was employed to perform whole-genome LTR-RT annotation and calculate the LTR Assembly Index (LAI) to assess the assembly’s continuity. The accuracy of the final assembly was also estimated from mapped *k*-mers via Merqury^19^. Illumina short reads were mapped back to the *de novo* assembly to identify potential errors such as SNPs or small insertions and deletions (indels). Additionally, the integrity of the *A. cristatum* genome assembly was verified by aligning PacBio HiFi reads (aligned using minimap2 v.2.21^63^) and Hi-C reads (processed with HICUP^60^) to the final assembly.

#### Genome annotation

#### Transcriptome sequencing

Total RNA was extracted from the stems, roots, leaves, stamens, pistils, and young spikes of *A. cristatum* Z559 using TRIzol reagent (Invitrogen, Carlsbad, CA, USA). cDNA libraries were constructed according to the Illumina standard protocol and sequenced on the Illumina platform using the PE150 strategy, generating 63.54 Gb of data (**Supplementary Table 9**). To construct a comprehensive catalog of transcript isoforms, equal amounts of total RNA from each tissue were pooled into a single sample for PacBio library preparation. Library construction and sequencing were performed according to the PacBio Iso-Seq protocol. Non-size-selected RNA from the pooled sample was run on the PacBio Sequel platform. Raw full-length cDNA reads were processed using the IsoSeq3 workflow (https://github.com/PacificBiosciences/IsoSeq), which involved trimming and clustering to generate high-quality transcriptome data. This process yielded 70.1 Gb of full-length transcriptome sequences (**Supplementary Table 9**). Additionally, other RNA-Seq data and full-length transcriptome sequences for *A. cristatum* Z559 were downloaded from the NCBI BioProject under accession number PRJNA534411^64^ and incorporated into this study. Raw sequencing reads were processed using Trimmomatic to remove low-quality reads and overrepresented sequences. Clean reads were aligned to the *A. cristatum* reference genome using HISAT2^65^. Only uniquely mapped reads with a perfect match or a single mismatch were retained for further analysis and annotation based on the reference genome. Gene expression levels were quantified in fragments per kilobase of transcript per million mapped reads (FPKM). Expression breadth was calculated using data from the five tissues mentioned above, considering only genes expressed in at least one tissue. Following the methodology described by Choulet et al.^66^, an FPKM threshold of 0.4 was used as the criterion for determining gene expression.

#### Protein-coding gene prediction and functional annotation

To predict and annotate protein-coding genes in the *A. cristatum* Z559 genome, we combined three approaches: *de novo* prediction, homology-based search, and transcript-based assembly. For *de novo* gene prediction, we utilized the ab initio software tools Augustus (v2.4)^67^ and SNAP (2006-07-28)^68^. Homology-based predictions were conducted using GeMoMa (v1.7)^69^ with reference gene models from several species, including *T*. *aestivum*^34^, *T. urartu*^53^, *T. turgidum*^23^, *Aegilops tauschii*^52^, *Hordeum vulgare*^24^, *Brachypodium distachyon*^70^, *Oryza sativa*^71^ and *Zea mays*^72^. Transcript-based predictions involved mapping RNA-seq data (described in transcriptome sequencing) to the reference genome using HISAT2 (v2.0.4), followed by assembly with StringTie (v1.2.3)^65^. Gene prediction from assembled transcripts was further refined with GeneMarkS-T (v5.1)^73^ and PASA (v2.0.2)^74^, which analyzed unigenes assembled by Trinity (v2.11)^75^. Gene models from all approaches were integrated using EVidenceModeler (EVM, v1.1.1)^74^ to produce a unified, non-redundant set of annotations. Evidence weights were assigned as follows: PASA-ISO-set > Homo-set > PASA-T-set > Cufflinks-set > Augustus > GeneID = SNAP = GlimmerHMM = Genscan. To ensure reliability, the models were filtered to exclude coding sequences (CDS) shorter than 300 bp, those lacking open reading frames, or those with premature stop codons.

Functional annotation of all protein-coding genes was performed by aligning them to multiple databases (**Supplementary Table 10**), including GenBank Non-Redundant (NR, 20200921)^76^, TrEMBL (202005), Pfam (33.1), SwissProt (202005)^77^, eukaryotic orthologous groups (KOG, 20110125), GO (20200615), and KEGG (20191220)^78^, using BLASTP (E-value < 1e-05). Additional plant TO annotations were sourced from NBRP and WheatOmics databases. Orthologous relationships, determined with Orthofinder (v2.3.1)^79^, were used to infer annotations for *A. cristatum* Z559 based on experimentally validated orthologs.

The resulting protein-coding genes were classified into HC and low-confidence categories. HC genes were supported by homology (≥80% identity and ≥50% coverage with HC gene sets of *T. urartu*, *Ae. tauschii*, *H. vulgare*, and *T. aestivum*) or transcriptome data (FPKM > 1) and lacked TE sequences. Low-confidence genes lacked homology or transcriptome support. The identification of alleles in the *A. cristatum* Z559 genome followed established methodologies^18^. The average copy number per gene is the total number of genes on the chromosome divided by the number of alleles. The average number of distinct alleles per gene is the total number of distinct coding sequence genes on the chromosome divided by the number of alleles.

#### TE annotation

The TEs in the *A. cristatum* genome were annotated using a combination of homology-based prediction and *de novo* methods. A nonredundant TE library was constructed by integrating *de novo* TE sequences, a wheat TE sequence library (ClariTeRep: https://github.com/jdaron/CLARI-TE), a plant TE sequence library (http://botserv2.uzh.ch/kelldata/trep-db/downloads/trep-db_complete_Rel-19.fasta.gz), and known databases, including Repbase (version 19.06), REXdb (version 3.0), and Dfam (version 3.2). TE sequences in the *A. cristatum* genome were identified and classified through homology searches against the combined library using RepeatMasker (http://www.repeatmasker.org).

Full-length LTR-RTs were identified using LTRharvest^80^ and LTR_finder^81^. High-quality intact LTR-RTs and a nonredundant LTR library specific to *A. cristatum* were generated using LTR_retriever^21^. In the LAI evaluation analysis, LTR_retriever was also used to determine the insertion times of intact LTR-RTs. The insertion time was calculated using the formula T = K/2μ, where μ represents a nucleotide substitution rate of 1.3 × 10^−8^ mutations per site per year, following previously established methods^23^. Additionally, the same annotation workflow was applied to other genomes for comparative analysis.

#### Cytological analysis

To detect *A. cristatum* genome chromatin in wheat–*A. cristatum* derivatives, genomic in situ hybridization (GISH) was performed on root tip cells using *A. cristatum* total genomic DNA labeled with Texas Red-5-dCTP (PerkinElmer, Boston, MA), and Fukuho genomic DNA was used as a blocker^82^. Nondenaturing FISH was carried out with oligonucleotide probes (oligo-AcCR2) to identify potential centromeric regions in *A. cristatum* Z559. The FISH probes were synthesized by Shanghai Sangon Biotech Co., Ltd. (Shanghai, China). All slides were examined using a Zeiss Axio Imager Z2 upright epifluorescence microscope (Carl Zeiss Ltd, Germany), with filters suitable for Vectashield mounting medium containing 4’,6-diamidino-2-phenylindole (DAPI). Hybridization signals were captured using a MetaSystems CoolCube 1 m CCD camera, and image analysis was performed with Metafer (automated metaphase image capture) and ISIS (image processing) software (MetaSystems GmbH, Germany).

#### ChIP-seq

ChIP-seq was performed on *A. cristatum* Z559 to identify centromeric regions, using a wheat-specific CENH3 antibody, with minor modifications to previously established protocols^83^. A custom anti-rabbit polyclonal antibody was designed against the N-terminal peptide sequence of *T. aestivum* TaαCENH3, “CARTKHPAVRKTK”. The purified antibody was dissolved in PBS buffer (pH 7.4) at a final concentration of 198 ng/μl. Nuclei were isolated from 2-week-old *A. cristatum* seedlings, digested with micrococcal nuclease, and incubated overnight at 4°C with either 3 μg of the CENH3 antibody or rabbit serum (negative control). Immune complexes were captured using Dynabeads Protein G, and chromatin was eluted with 100 μl of preheated (65°C) elution buffer containing 1% sodium dodecyl sulfate and 0.1 M NaHCOL. DNA was subsequently isolated using the ChIP DNA Clean & Concentrator Kit, and ChIP-seq libraries were prepared with the TruSeq ChIP Library Preparation Kit. High-quality 150-bp paired-end reads were generated on the NovaSeq S4 platform.

The ChIP-seq reads were aligned to the reference genome using BWA^84^, and only reads with a mapping quality score of 30 or higher were retained to ensure data reliability and filter out non-unique sequences. Centromere boundaries were defined by identifying peaks of CENH3 enrichment using the epic2 peak caller^85^. These boundaries were further refined by analyzing the density of epic2 peaks at a resolution of 500 kb.

#### Circos plot

The distributions of genes, transcripts, TEs, expression breadth, and CENH3 ChIP-seq coverage were visualized in circos plot within a sliding window of 20 Mb with a step size of 5 Mb. Synteny blocks between the four haplotypes were identified using MCScanX^86^. The circos plot was generated with the *circlize*^87^.

### Comparative genomic analysis

#### Gene-level synteny between *A. cristatum* and other Triticeae species

A process similar to that described by Choulet et al.^66^ was employed to filter HC genes in *A. cristatum* Z559, wheat (cv. Chinese Spring), rye (cv. Lo7), and barley (cv. Morex), generating core gene sets for comparative analyses. In this process, TEs and alternative splice variants were removed from the gene sets of all species. The protein sequences of filtered HC genes from *A. cristatum*, wheat, rye, and *Th. elongatum* were reciprocally aligned using BLASTP with an E-value threshold of < 1e-05. A collinear block was defined as a conserved set of at least five genes (anchors) in the same order between two genomes, with a maximum of 25 spacer genes between the anchors within a block. The reciprocal best-hit alignment for each pair was used to construct whole-genome collinearity between *A. cristatum*, rye, *Th. elongatum*, and *T. aestivum* using MCScanX^86^ with default parameters. The jcvi software (https://github.com/tanghaibao/jcvi/wiki) was employed to visualize synteny results, detect large structural variations and collinearity blocks between *A. cristatum*, rye, *Th. elongatum*, and *T. aestivum*.

#### Phylogeny and divergence time analysis

Protein sequences from *A. cristatum* and 13 other plant species, including *Setaria italica*^88^, *Zea mays*, *Sorghum bicolor*^89^, *Oryza sativa*, *Brachypodium distachyon*, *Hordeum vulgare*, *Secale cereale*, *Th. elongatum*, *T. urartu*, *T. aestivum*, and *Ae. tauschii*, were used for gene family construction. Sequences shorter than 30 amino acids were filtered out, and protein clustering was performed using CD-HIT-EST^90^ to select the longest sequences from highly similar clusters for each species. The protein sequences for all plant species were grouped into orthologous genes using OrthoFinder with the following parameters: -S blast -t 70 -M msa -A muscle -T raxml-ng -I 1.5. The protein sequences of the deduced single-copy genes were aligned using MAFFT^91^ for multiple sequence alignment, and conserved sequences were extracted using trimAl^92^. A phylogenetic tree was constructed by BEAST^93^ based on the Gamma site model using the GTR+G substitution model with 1,000 bootstrap replicates.

To identify and analyze collinearity anchor pairs, we used the WGDI package^94^. First, all syntenic blocks were identified using the improved collinearity pipeline in WGDI with a significance threshold of “P value = 0.05”. The *Ks* value for each anchor gene pair located within these syntenic blocks was calculated using the *Ks* pipeline in WGDI. A *Ks* dot plot of all anchor pairs was generated using the block pipeline, and the kspeaks pipeline was applied to analyze the distribution of the *Ks* median value for each syntenic block. Finally, the results of these analyses were summarized in a single figure using the ggplot2 package. The divergence and WGD occurrence times were calculated using the formula *K*sL=L*t*/2*r*, where *t* is the time of divergence and *r* is the mutation rate.

#### Specific and expanding gene families

Gene families specific to *A. cristatum* were identified based on the results of Orthofinder analysis. To examine the evolution of gene family sizes, the CAFÉ v3.1 software^95^ was used, which is based on the stochastic birth-death model. Using the previously calculated phylogeny and divergence times, CAFÉ was applied to detect gene families that have undergone expansion in *A. cristatum* and the other species analyzed, with the parameters “p-value = 0.05, number of random simulations = 1000, and search for lambda”.

#### Positive selection sites

To identify positive selection sites, CDS alignments for single-copy gene families from *A. cristatum* Z559 genome, *T. urartu*, *Ae. tauschii*, *T. aestivum* subgenomes, *H. vulgare*, *T. elongatum*, *S. cereale*, *B. distachyon*, *O. sativa*, *S. bicolor*, and *S. italica* were generated using MUSCLE^96^. Poorly aligned positions and divergent regions were filtered using Gblocks^97^. Positive selection sites in *A. cristatum* were identified using the branch-site models in PAML^98^, with *A. cristatum* as the foreground branch. P-values were calculated using the χ^2^ statistic and adjusted for false discovery rate (FDR).

#### GO, KEGG and TO enrichment analysis

Hypergeometric tests were performed to assess whether specific functional categories from GO, Plant TO^99^, and KEGG^78^ were significantly overrepresented in different gene sets of *A. cristatum* within the genome. Functional enrichment analyses were conducted using the R package clusterProfiler (version 4.0)^100^, with the background set consisting of all annotated genes in *A. cristatum* in this study.

#### Characterization of *A. cristatum* introgressions in wheat-*A. cristatum* derivatives

We generated Illumina short-read sequencing data ranging from approximately 0.5 × to 10 × for 134 wheat–*A. cristatum* addition and translocation lines. The cleaned sequencing reads were aligned to a pseudo-genome (ABDP_1), constructed by concatenating the reference genomes of wheat cultivar ‘Chinese Spring’ and the longest haplotype genome of *A. cristatum* Z559. PCR duplicates were identified and removed using Picard’s MarkDuplicates tool (https://broadinstitute.github.io/picard/). Alignments were filtered based on the following criteria: 1) properly paired reads on chromosome P, 2) alignment score (SAM Tag AS:i) > 90, 3) edit distance to the reference (SAM Tag NM:i) < 13, and 4) mapping quality (MAPQ) > 30. Introgression regions were determined by comparing the coverage distribution with that of the wheat parent, Fukuho.

In contrast, the 297 wheat–*A. cristatum* breeding derivatives, developed through multiple hybridizations, have complex genetic backgrounds, making it challenging to detect foreign gene introgressions using traditional alignment-based methods. Instead, we employed the *k*-mer-based IBSpy approach, which has proven effective for detecting alien introgressions^27,29^. We generated 10 × resequencing data for all breeding derivatives and their 10 wheat parental lines. To reduce false positives, we also downloaded resequencing data from 74 additional wheat varieties for this study. Using KMC (v3.1.2)^101^, we constructed 31-mer sequence sets for all resequenced lines. With the *A. cristatum* Z559 genome as the reference, we applied IBSpy with a *k*-mer size of 31 and a window size of 50,000 bp to analyze the variant counts across all samples. Introgression lines were identified by comparing their *k*-mer variant counts to those observed in the 10 parental lines and the 74 wheat varieties. If the number of variants in a specific genomic window of a breeding derivative was significantly lower than that of its parental lines and all wheat accessions (P value of the χ^2^ test < 0.05), the region was identified as an introgressed fragment.

#### Growth conditions of wheat-*A. cristatum* derivatives

To evaluate the field phenotypes of the tested materials, field trials were conducted during the 2021-2022 growing season at the experimental station of the Chinese Academy of Agricultural Sciences (Xinxiang, Henan, China). All wheat-*A. cristatum* derivatives were grown at this site. The wheat-*A. cristatum* addition lines del26 and del19a, along with the control wheat cultivar Fukuho, were sown in three single rows per material, each row measuring 2 m in length, with 20 seeds per row and a row spacing of 0.3 m. For the 141 wheat-*A. cristatum* breeding derivatives, each line was grown in a plot consisting of six rows, each 4.5 m long, with a row spacing of 0.25 m and 300 seeds per row, ensuring a planting density consistent with actual wheat production. Sowing and harvesting were fully mechanized, and grain yield was determined based on the total harvested weight per plot.

### Cloning of *AcGNS1* gene

#### Transcriptome sequencing of addition line del26

Total RNA was extracted from the mature stems, roots, leaves, and young spikes at the double ridges and spikelet differentiation stages of wheat Fukuho-*A. cristatum* addition line del26 using TRIzol (Invitrogen, Carlsbad, California). cDNA libraries were constructed following the Illumina standard protocol and sequenced on the Illumina platform using the PE 150 strategy. Raw sequencing data were processed using Trimmomatic to remove low-quality reads and overrepresented sequences. The clean reads were then mapped to the pseudo-genome ABDP_1 using HISAT2^65^. Only uniquely mapped reads with perfect matches were further analyzed. Gene expression levels were estimated by FPKM.

#### Real time PCR analysis

Total RNA was extracted using a plant total RNA extraction kit (ZOMANBIO) according to the manufacturer’s instructions. RNA integrity was analyzed by spectrophotometry and 1% agarose gel electrophoresis. Reverse transcription was performed using a kit (ZOMANBIO). The complementary DNA (cDNA) was subjected to real-time RT-PCR using a kit (Takara) in a Step One Plus Real-Time PCR System (Applied Biosystems) and detected using software. Real-time RT-PCR was performed with six biological replicates, and wheat *TaActin* was used as a reference gene. Relative gene expression data were analyzed using the 2^−^(ΔΔCt) method.

#### Generation of transgenic wheat plants

To generate the overexpression construct, the CDS of *AcGNS1* was cloned from the wheat-*A. cristatum* partial chromosome addition line del26 into the pCAMBIA3300 vector (using BamHI/SmaI), driven by the maize Ubiquitin promoter, using an In-Fusion HD Cloning Kit (Takara). The constructs were then transformed into *Agrobacterium tumefaciens* strain GV3101, which was used to transform wheat callus by the *Agrobacterium*-mediated transformation method. The wheat variety Fielder was selected as the receptor. Three homozygous T_3_ lines of *AcGNS1* transgenic plants were used in this study. All transgenic lines and the recipient wheat cultivar Fielder were grown in the transgenic experimental fields in Shunyi, Beijing. Each line was sown in three single rows, with each row measuring 2 m in length, containing 20 seeds, and spaced 0.3 m apart. Conventional water and fertilizer management practices were applied throughout the growing period.

#### Heritability assessment and building GS models

SNPs were also determined for 297 wheat-*A. cristatum* breeding derivatives following the GATK variant calling pipeline (https://gatk.broadinstitute.org/). Phenotypic values for 6 yield-related traits were also determined for 141 of the 297 breeding derivatives during 2021-2022. Heritability was estimated as the proportion of phenotypic variance explained by only SNPs and by SNPs and introgression segments using the LDAK model^102^.

GS models were then built for each of the six yield-related traits in 141 breeding derivatives following the CropGBM pipeline^103^. In short, genotypic and phenotypic data were split into training (60%), validating (20%) and testing (20%) datasets. GS models were trained using training and validating datasets. On average, for each trait, the top 700 to 800 trait-associated markers were selected from CropGBM pipeline for phenotype prediction. Model prediction accuracies were determined by evaluating models using hold-out test datasets. Both SNP-only model and SNP+introgression model were built and evaluated. GEBVs were estimated using built models for all 297 breeding derivatives that have been resequenced and genotyped. The line with the highest ranked yield GEBV, Pubingzi 300, was subsequently selected for national regional trials.

#### Regional trial of new wheat varieties in China

In the 2022-2023 and 2023-2024 growing seasons, trials for the candidate wheat variety Pubingzi 300 were conducted at 23 and 17 sites, respectively, across key wheat-producing provinces in the Southern Huang-Huai region of China, including Henan, Anhui, Jiangsu, and Shaanxi (**Supplementary Table 26**). All trials were carried out in local production fields to reflect realistic agronomic conditions. The experimental design followed a sequential arrangement without replication, with a minimum plot size of 0.033 hectares. Sowing and harvesting were fully mechanized, and yields were determined based on the total plot weight.

To adapt to regional agronomic practices, the target seedling density at each site was set according to the recommended guidelines for the predominant local wheat variety, ensuring uniformity across all trial plots within a site. Sowing was completed on a single day at each site, strictly adhering to the recommended seeding rate and sowing period to minimize environmental variability. Zhoumai 36, a widely grown wheat variety, was used as the standard control across all locations.

Field management was optimized to reflect high-input local agronomic practices, with adjustments made according to the specific ecological conditions of each trial site. This comprehensive trial design aimed to assess the adaptability, yield potential, and agronomic performance of the new wheat variety Pubingzi 300 under realistic field conditions.

## Data availability

The raw sequencing data used for de novo genome assemblies, the RNA-seq and Iso-seq data for the annotation, the ChIP–seq reads and the whole-genome sequencing reads of 431 wheat-*A. cristatum* introgressions are available at the National Center for Biotechnology Information (NCBI) SRA database under BioProject accession PRJNA906942. The genome assembly data have been deposited in the NCBI database under BioProject accession PRJNA1281305, and in the Chinese National Genomics Data Center (https://bigd.big.ac.cn/) under BioProject accession PRJCA042075.

## Acknowledgements

This publication is based on work supported by the National Natural Science Foundation of China (32472107 to S.Z.), the National Key Research and Development Program of China (2021YFD1200605 to L.L.), the Basic Research Center, Innovation Program of Chinese Academy of Agricultural Sciences (CAAS-BRC-CS-2025-01 to W.Q.).

## Contributions

L.L. managed the project. L.L., S.Z., Q.H., and H.-Q.L. conceived and designed the research. S.Z., H.H., J.Z., and X.Y. maintained and provided plant materials. S.Z. performed genome sequencing, assembly, and validation; characterized the tetraploid *A. cristatum* genome; conducted genome evolution analysis, TE analysis, ChIP-seq, comparative genomics, and centromere analysis; and analyzed wheat–*A. cristatum* addition and translocation lines. S.Z., J.-S.Z., and B.G. conducted genome annotation, gene expression analysis, and gene functional enrichment analysis. B.G. identified key gene copy numbers and homologs of known functional genes in the addition lines. S.Z., G.B., B.H., and H.H. collected RNA-seq and genome resequencing samples. W.Y. cloned the *AcGN S1* gene and performed phenotypic analysis of transgenic materials. H.H. and B.H. performed cytological experiments. L.L. developed the wheat–*A. cristatum* breeding derivatives. S.Z., B.G., B.H., X.L., Y.-D.L., and W.Y. conducted phenotypic evaluations. P.Z. and Q.H. performed genotyping of wheat–*A. cristatum* breeding derivatives, analyzed *A. cristatum* introgressions into wheat, conducted heritability assessments, and constructed GS models. Y.D. evaluated the genome QV. Y.D., B.H., and B.L. contributed to figure preparation. J.Z. conducted regional trials of the new wheat varieties. S.Z., Q.H., L.L., and H.-Q.L. wrote the manuscript with input from N.S., Z.L., Y.X., P.Y., K.Y., J.Z., and Y.L. All authors have read, edited, and approved the content of the manuscript.

**Extended Data Table 1.** Summary of genome assembly and annotation of *A. cristatum* Z559

| Genomic feature | *A. cristatum* Z559 | Haplotype 1 | Haplotype 2 | Haplotype 3 | Haplotype 4 |
| --- | --- | --- | --- | --- | --- |
| Estimated genome size (Gb) by K-mer | 24.4 |  |  |  |  |
| Assembled genome size (Gb) | 25.62 | 6.33 | 6.11 | 6.02 | 5.91 |
| Percent of assembled genome size (%) | 100 | 24.71 | 23.85 | 23.5 | 23.07 |
| Anchored genome size (Gb) | 24.37 |  |  |  |  |
| Percent of anchored genome (%) | 95.12 |  |  |  |  |
| Contig N50 (Mb) | 26.22 | 28.04 | 33.4 | 25.19 | 27.15 |
| Scaffold N50 (Mb) | 876.63 | 887.7 | 851.5 | 831.3 | 818.54 |
| BUSCO completeness of assembly (%) | 98.7 | 97.7 | 97.4 | 97.3 | 96.5 |
| LAI score | 22.96 |  |  |  |  |
| QV | 41.81 |  |  |  |  |
| Total number of genes | 171,519 | 41,160 | 40,061 | 40,141 | 39,813 |
| BUSCO completeness of annotation (%) | 97.5 |  |  |  |  |
| Size of retrotransposons (Gb) | 20.01 | 5 | 4.81 | 4.75 | 4.64 |
| Percent of retrotransposons (%) | 78.1 | 78.99 | 78.72 | 78.9 | 78.51 |
| Size of DNA transposons (Gb) | 2.63 | 0.6 | 0.58 | 0.57 | 0.56 |
| Percent of DNA transposons (%) | 10.27 | 9.48 | 9.49 | 9.47 | 9.48 |
| Size of total repeat (Gb) | 22.24 | 5.59 | 5.38 | 5.3 | 5.2 |
| Percent of repeat (%) | 86.81 | 88.31 | 88.05 | 88.04 | 87.99 |

**Extended Data Fig. 1.**
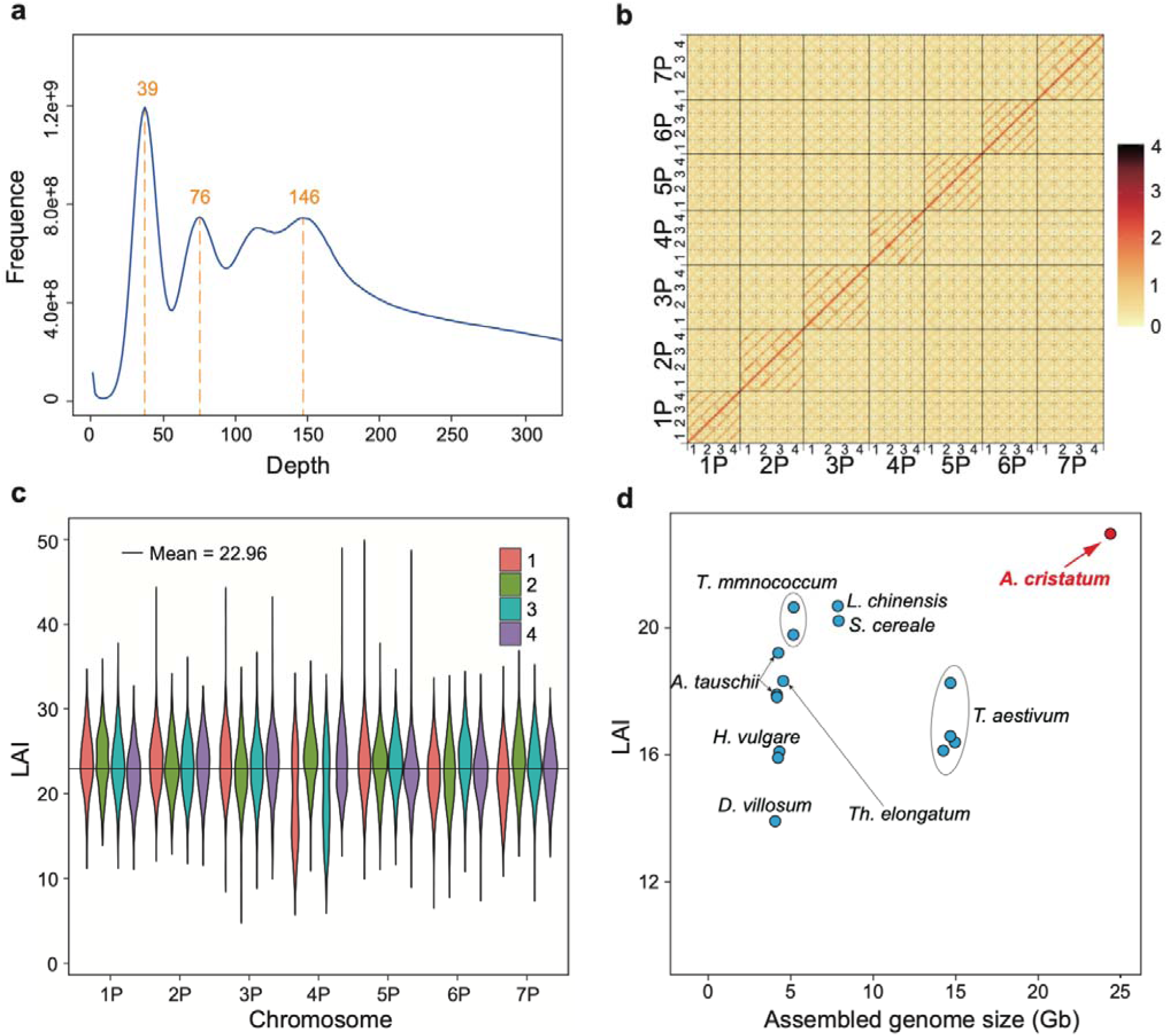
Evaluation of the assembled *A. cristatum* Z559 genome. **a,** *K*-mer distribution used to estimate the genome size and heterozygosity of the *A. cristatum* Z559 genome. The genome size was calculated by dividing the total *K*-mer count by the coverage depth (frequency peak at 39). The secondary peaks at 76 and 146 correspond to high heterozygosity and autotetraploidy, respectively. **b,** Hi-C interaction heatmap at 500-kb resolution. Intrachromosomal interactions are visualized as antidiagonal patterns, demonstrating the chromosomal architecture of the assembly. **c,** Boxplot of the LTR Assembly Index (LAI) distribution for each chromosome. The average LAI of *A. cristatum* Z559 is approximately 22.96 (red line), exceeding the gold-standard quality threshold of LAI = 20 and confirming the high quality of the assembly. **d,** Reported genome sizes and LAI scores of representative Triticeae genome assemblies. *A. cristatum* Z559 (this study) is highlighted in red, emphasizing its high assembly quality and the largest genome size compared to other Triticeae genomes.

**Extended Data Fig. 2.**
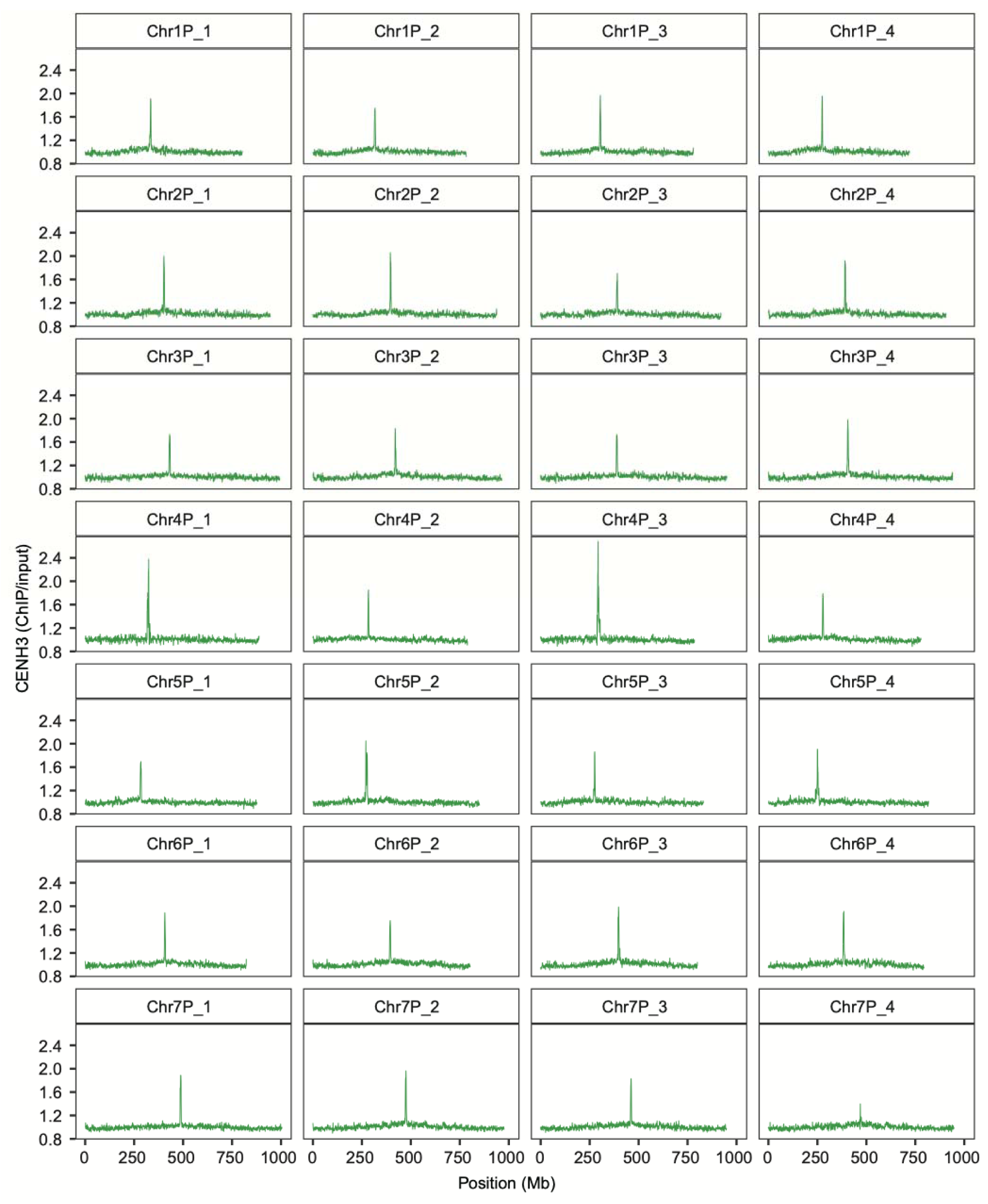
CENH3 ChIP-seq read coverage normalized by input across all chromosomes of *A. cristatum* Z559. CENH3 ChIP-seq read coverage was normalized using input control data across all chromosomes. The ChIP/input ratio was calculated with bamCompare at a resolution of 500-kb bins.

**Extended Data Fig. 3.**
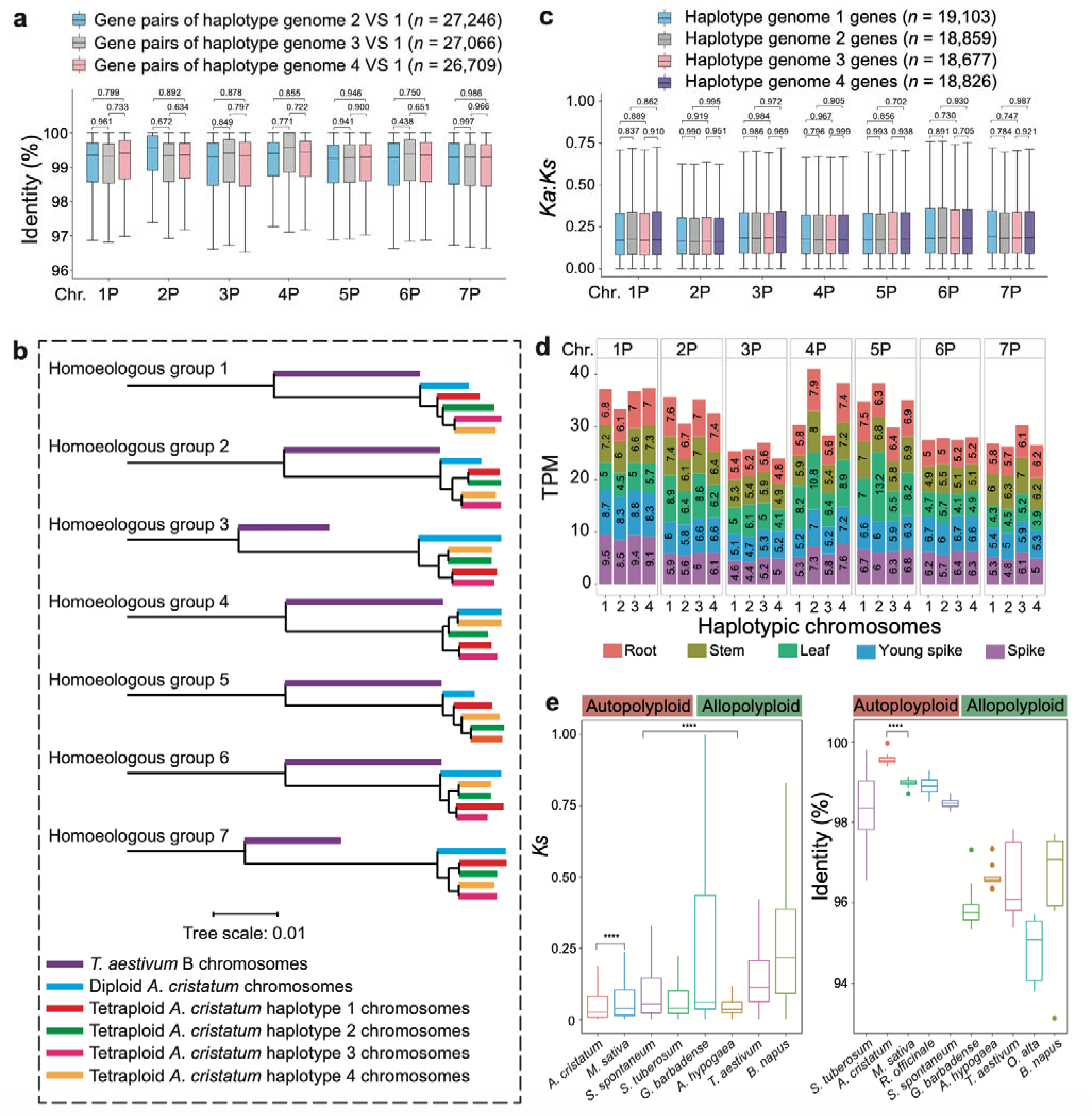
Comparative analysis of haplotypic chromosomes of *A. cristatum* Z559. **a,** Boxplots illustrating the identity distributions of homologous genes among chromosomes with 2, 3, or 4 haplotypes compared to 1 haplotype chromosomes in *A. cristatum* Z559 (excluding genes in scaffolds). **b,** The phylogenetic tree topology constructed based on single-copy orthologous genes between the haplotypic chromosomes of *A. cristatum* Z559 and the chromosomes of a diploid *A. cristatum* genome (unpublished). The wheat B subgenome was used as the outgroup. **c,** Boxplots of the *Ka*:*Ks* ratio distributions for homologous genes from each tetraploid chromosome of *A. cristatum* Z559, with the *T. aestivum* D genome as a reference. Genes on scaffolds or lacking *Ka*:*Ks* values were excluded. Blue, gray, pink, and purple boxplots represent *Ka*:*Ks* distributions for each tetraploid group. **d,** Total allelic expression from om *A. cristatum* Z559 chromosomes in mature leaves, roots, stems, spikes, and young spikes. Numbers 1–4 represent the four haplotypic chromosomes in each group. **e,** Distributions of *Ks* (left) and identity (right) for gene pairs among different monoploid genomes. a, c and e, The boxplots show the 25th, 50th, and 75th percentiles, with whiskers extending to 1.5× the interquartile range. *P* values were calculated using a two-sided t-test. *n* values in brackets indicate the number of genes analyzed in a and c.

**Extended Data Fig. 4.**
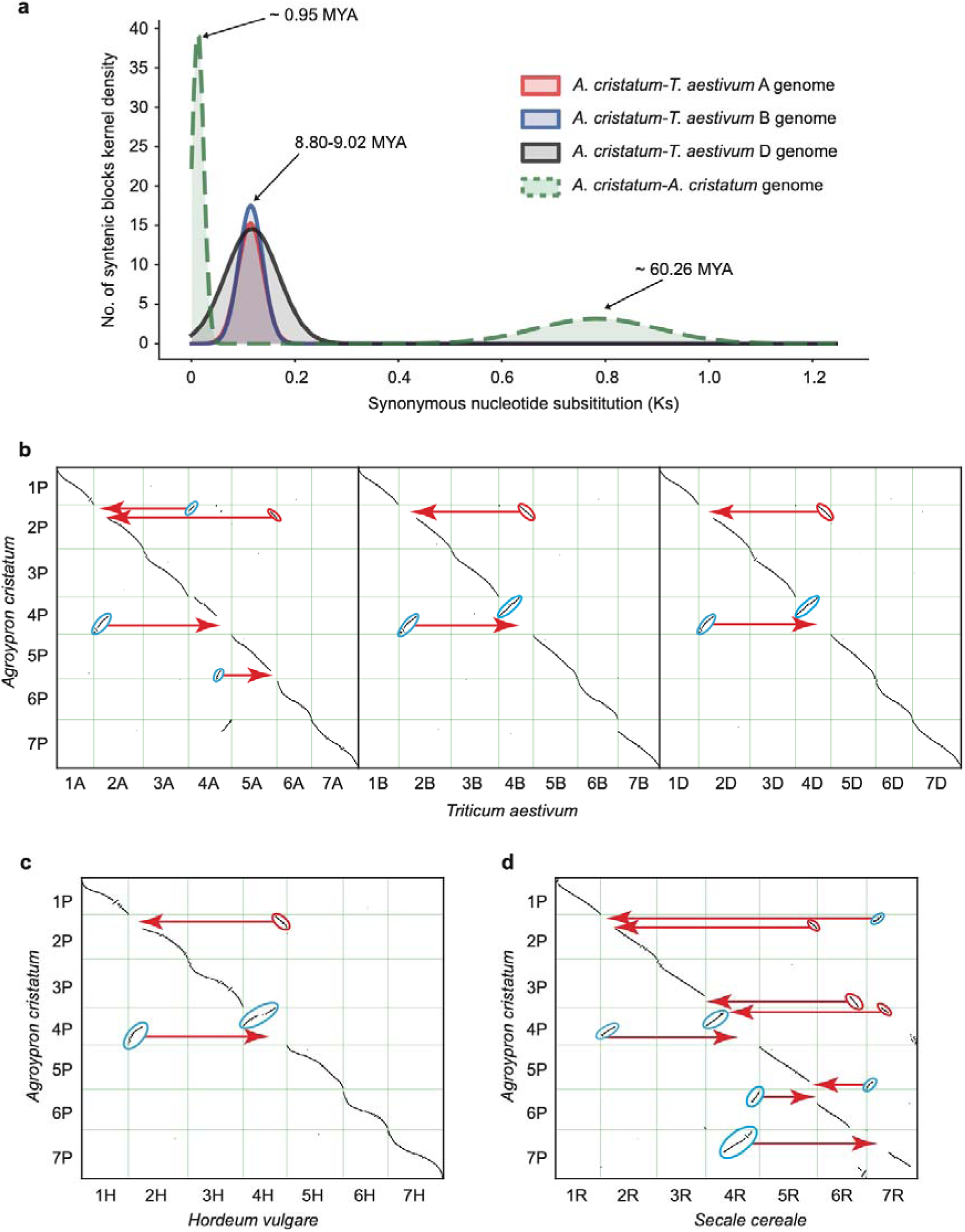
Comparative genomics of *A. cristatum*. **a,** The *Ks* distribution within the *A. cristatum* genome and in comparison with wheat subgenomes. Peaks in the *Ks* distribution and their corresponding divergence times are marked with arrows and labels. **b-d,** Gene collinearity between the *A. cristatum* genome and the genomes of *T. aestivum* (b), *H. vulgare* (c), and *S. cereale* (d). The vertical axis represents the positions of *A. cristatum* chromosomes, while the horizontal axis represents the chromosomes of *T. aestivum*, *H. vulgare*, and *S. cereale*. Rectangles indicate chromosomal

**Extended Data Fig. 5.**
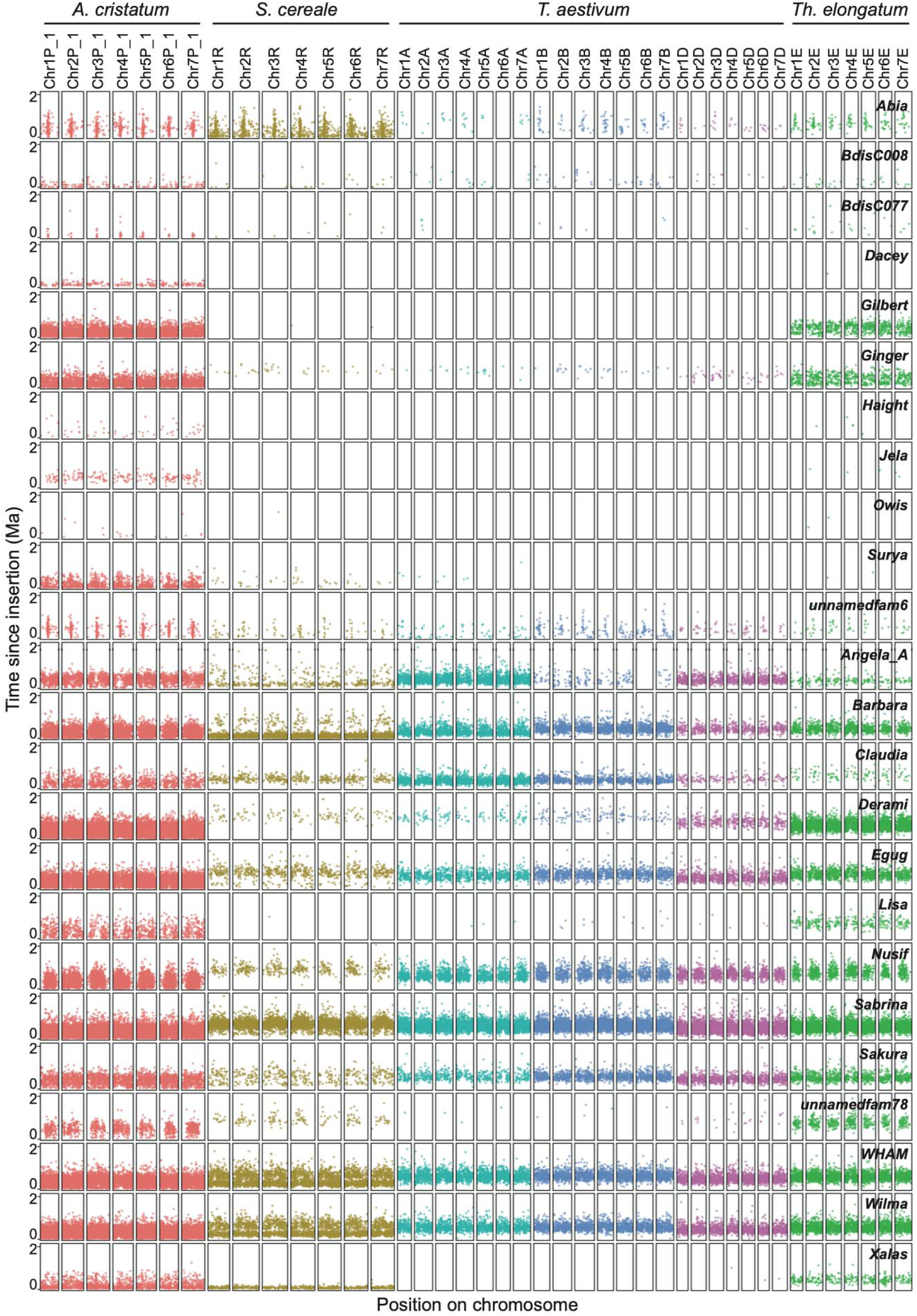
Abundance of full-length LTR-RT families in *A. cristatum* genome. Twenty-four full-length LTR-RT families show significantly elevated abundance in *A. cristatum* genome

**Extended Data Fig. 6.**
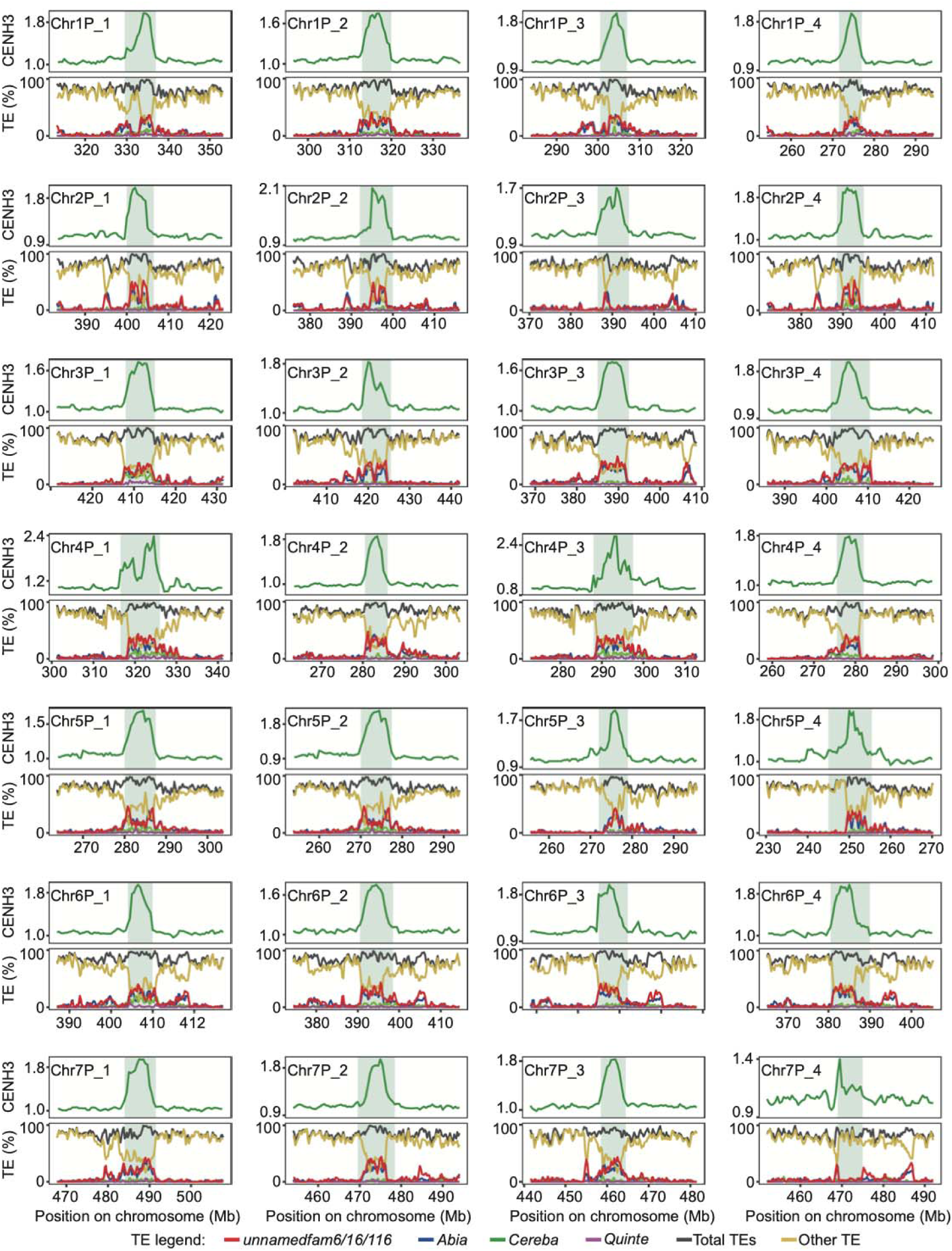
Identification of functional centromeres in *A. cristatum* Z559. Functional centromeres (shaded in blue) were defined using CENH3 ChIP-seq data. Centromeric regions and approximately 20 Mb of flanking regions are displayed. Top, the ratio of CENH3 ChIP-seq reads to input control reads. The functional centromeric regions were determined based on epic2 ChIP-seq peak call density, highlighted by the shaded areas. Bottom, average TE content in 100 kb windows. Notably, *unnamedfam6*/*16*/*116* and *Abia* retrotransposons are highly enriched in functional centromeres, whereas

**Extended Data Fig. 7.**
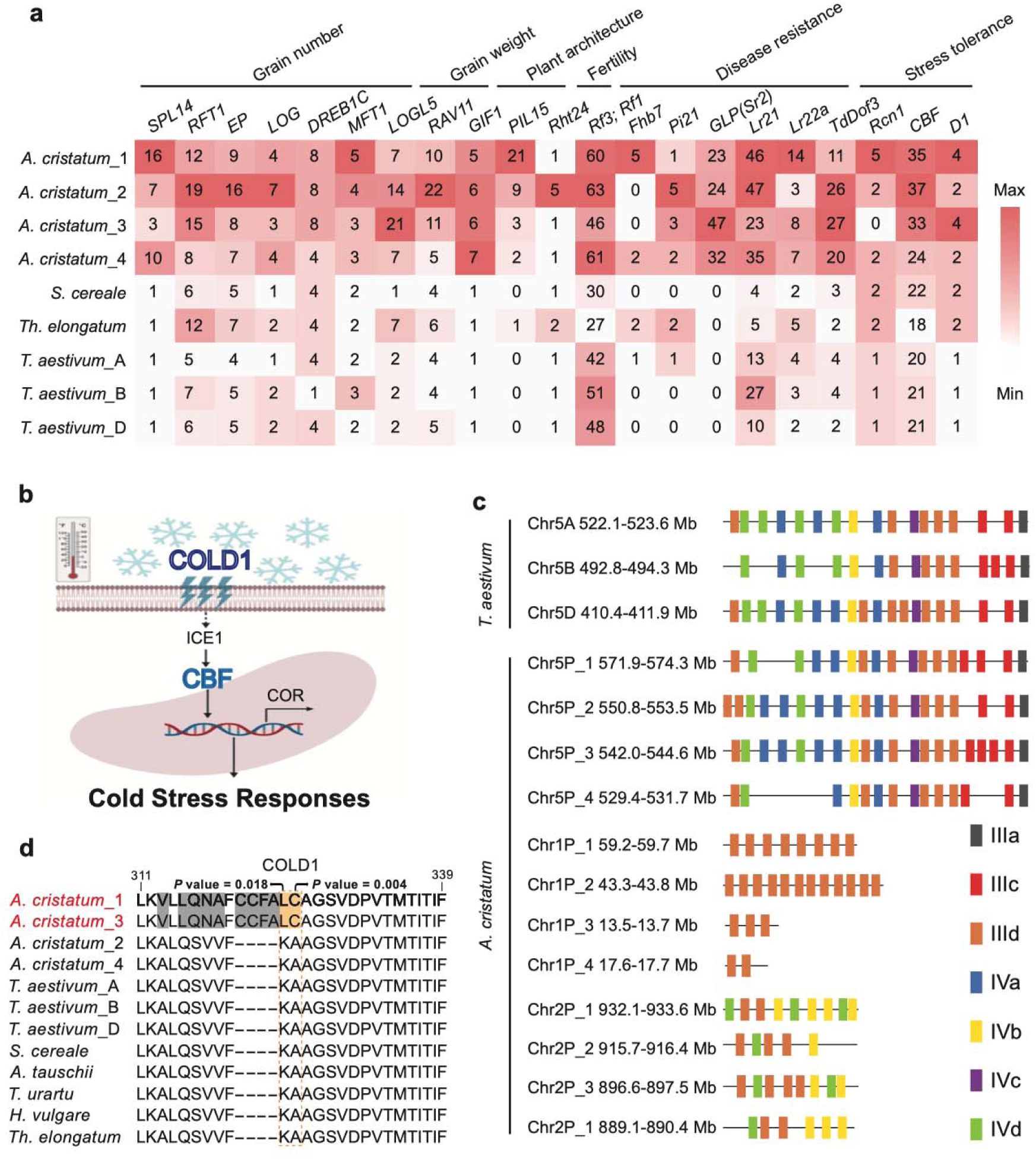
Specificity of genes in *A. cristatum* Z559 genome. **a,** Expanded vital functional gene families likely associated with agronomically important traits in the four haplotype genomes of *A. cristatum*, compared with rye (*S. cereale*), *Thinopyrum elongatum* (*Th. elongatum*), and wheat subgenomes. Gene family counts are displayed as a heat map. **b,** Illustration of cold sensing in plant cells. **c,** Genomic location of *CBF* gene duplication in *A. cristatum* compared to the corresponding region in wheat. The *CBF* gene cluster on homologous group 5 constitutes the *Fr2* locus, while two additional clusters on chromosomes 1P and 2P are specific to the P genome. **d,** The positively selected gene *COLD1* in the P genome. Sites showing variation between genomes are highlighted in different colors, with positively selected variation sites between the P genome and outgroups marked by dashed boxes.

**Extended Data Fig. 8.**
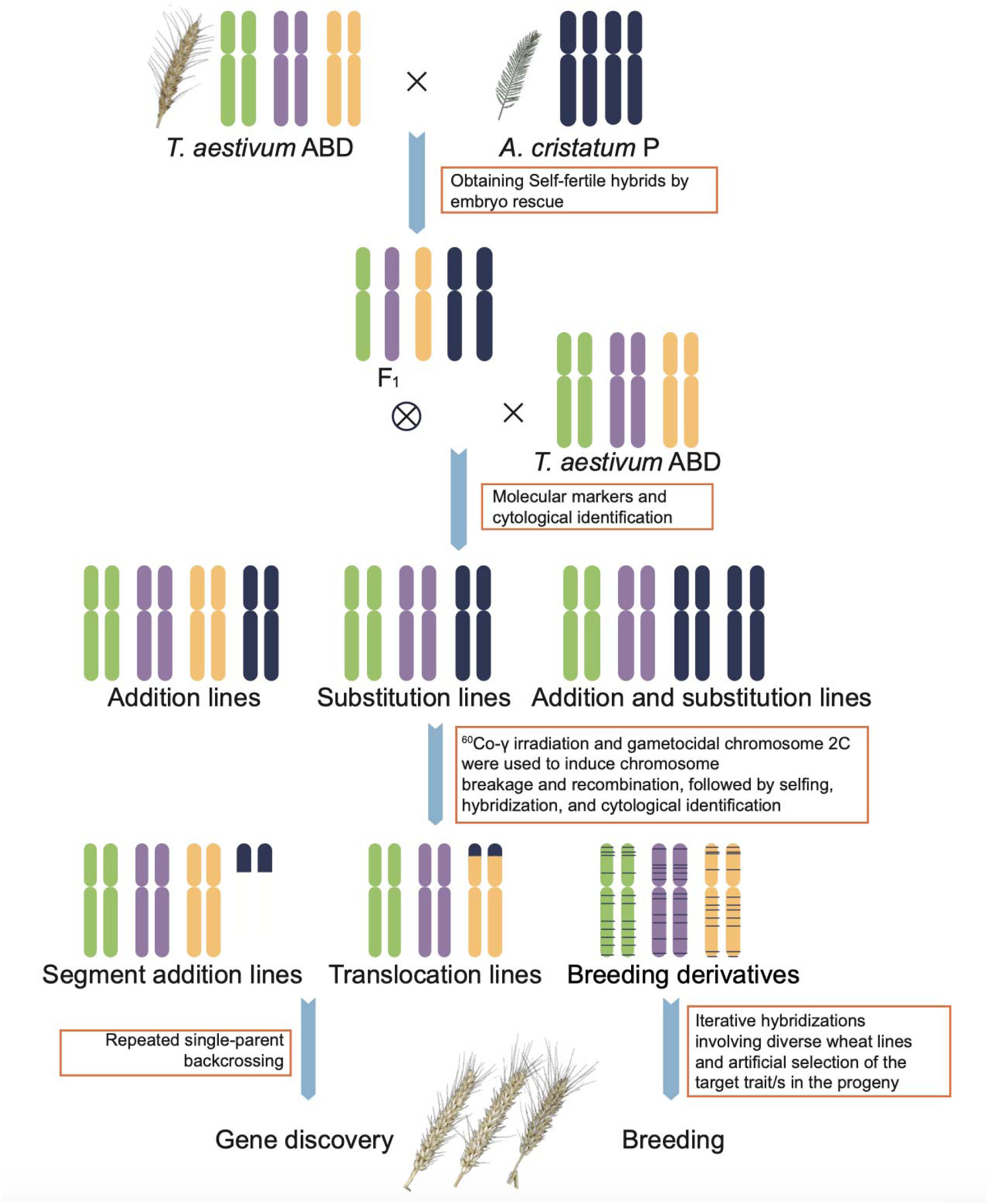
Flowchart illustrating the process of distant hybridization between wheat and *A. cristatum* and the subsequent generation of wheat-*A. cristatum* derivatives.

