## Supplementary Figures for "*Agropyron cristatum* genome provides a new insight for wheat improvement with wild relatives"

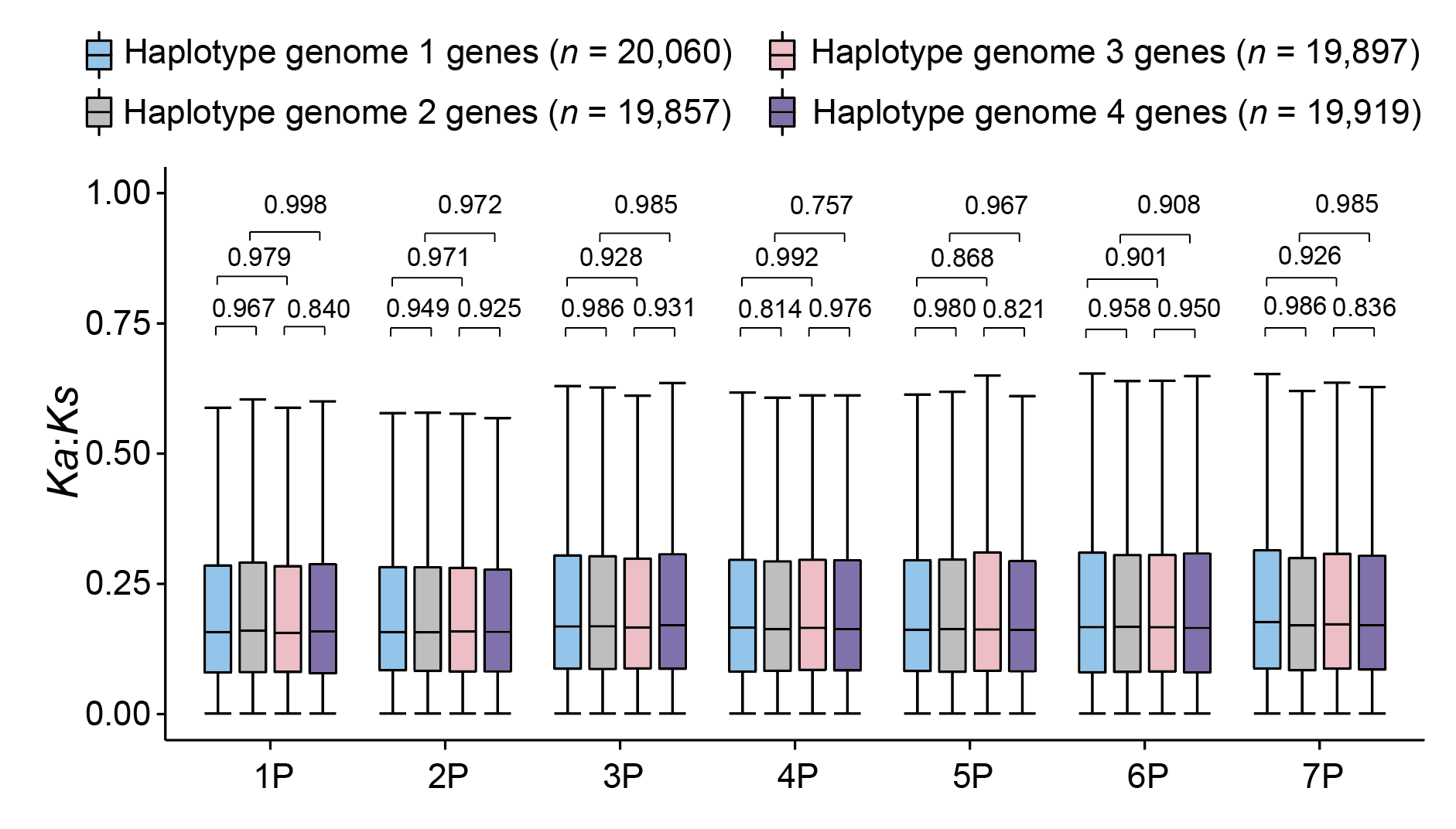
**Supplementary Fig. 1 Distribution of Ka:Ks ratios in homologs from A. cristatum tetraploids compared with Hv genome orthologs.** Boxplots represent the Ka:Ks ratio distributions for homologous genes from each tetraploid in the A. cristatum genome, using Hv genome orthologs as references (excluding genes on scaffolds or without Ka:Ks ratios). The blue, gray, pink, and purple boxplots indicate the Ka:Ks distributions of homologous genes from the four A. cristatum tetraploids. The boxplots display the 25th, 50th (median), and 75th percentiles, with whiskers extending to 1.5× the interquartile range. P values were determined using a two-sided t-test. The n values in brackets indicate the number of genes analyzed for each type on the chromosomes.

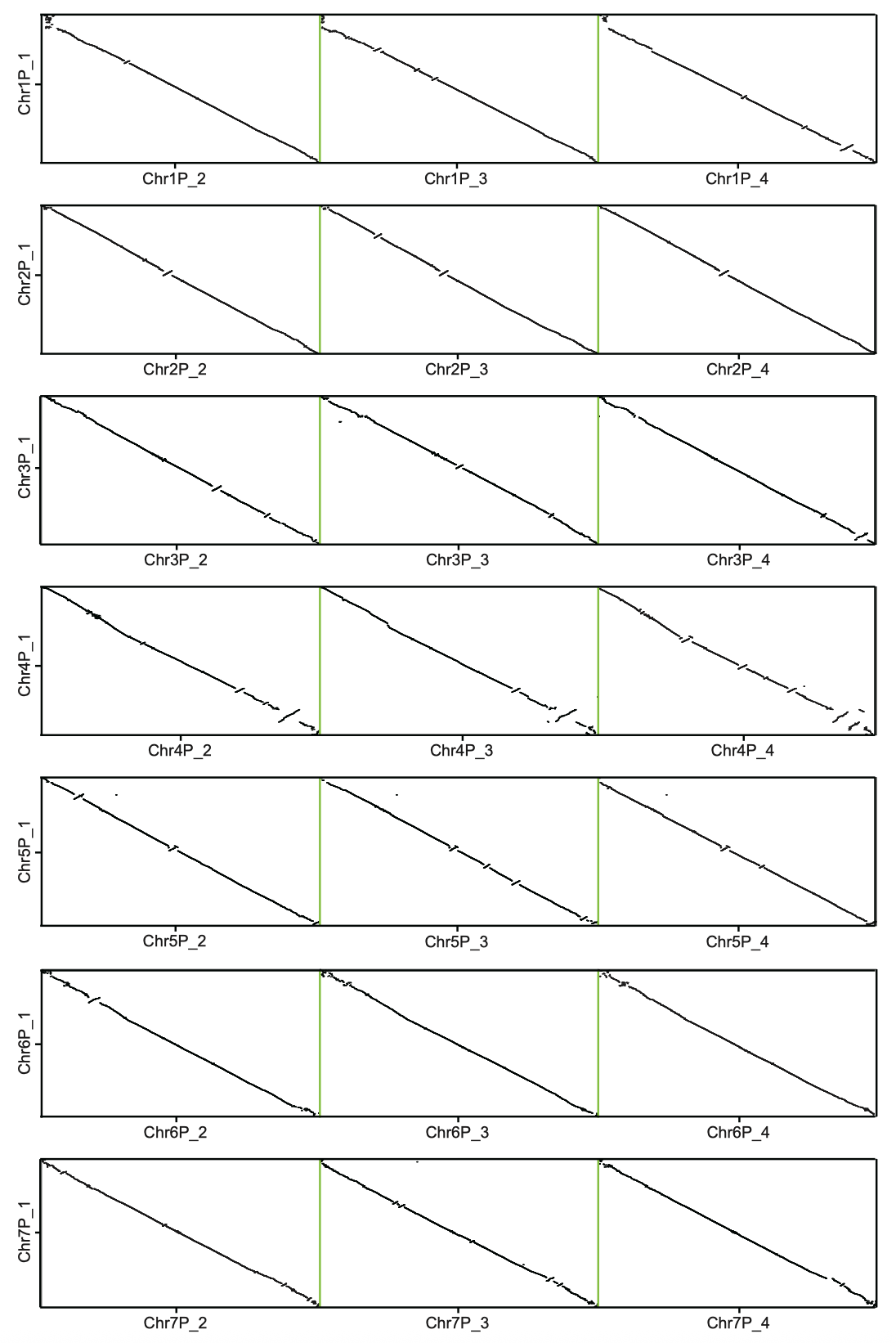
**Supplementary Fig. 2. Structural rearrangements between the four haplotypes of each chromosome.**

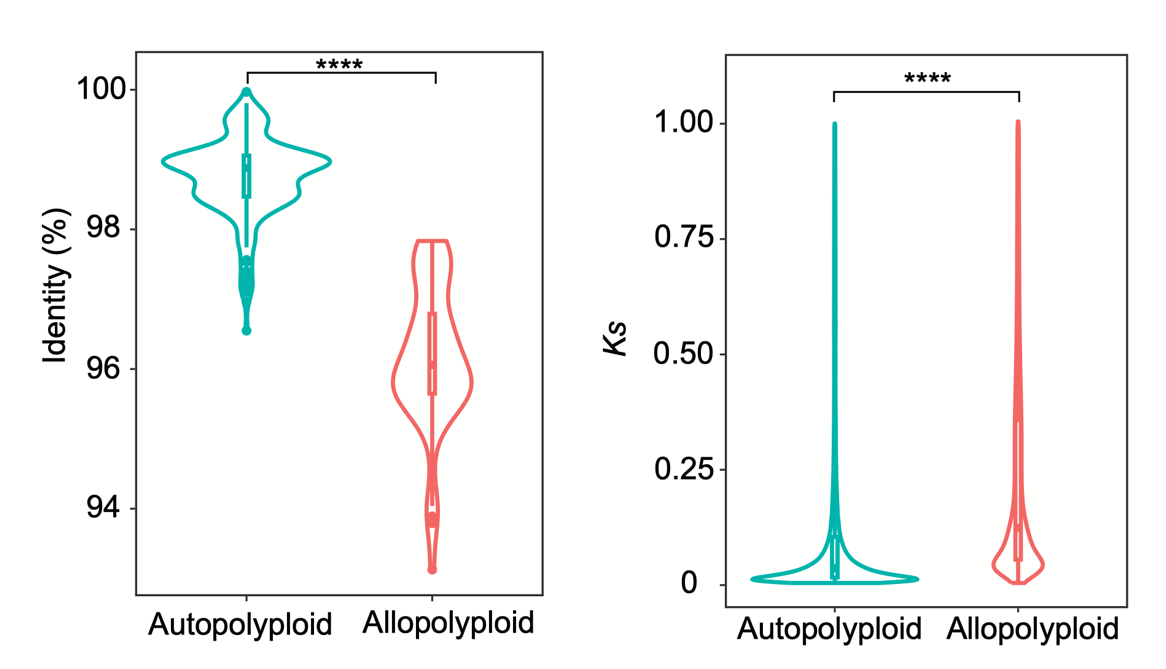

**Supplementary Fig. 3 Sequence identity and Ks of homologous chromosomes in autopolyploid and allopolyploid genomes.** Blue represents autopolyploids, and red represents allopolyploids (*t*-test).

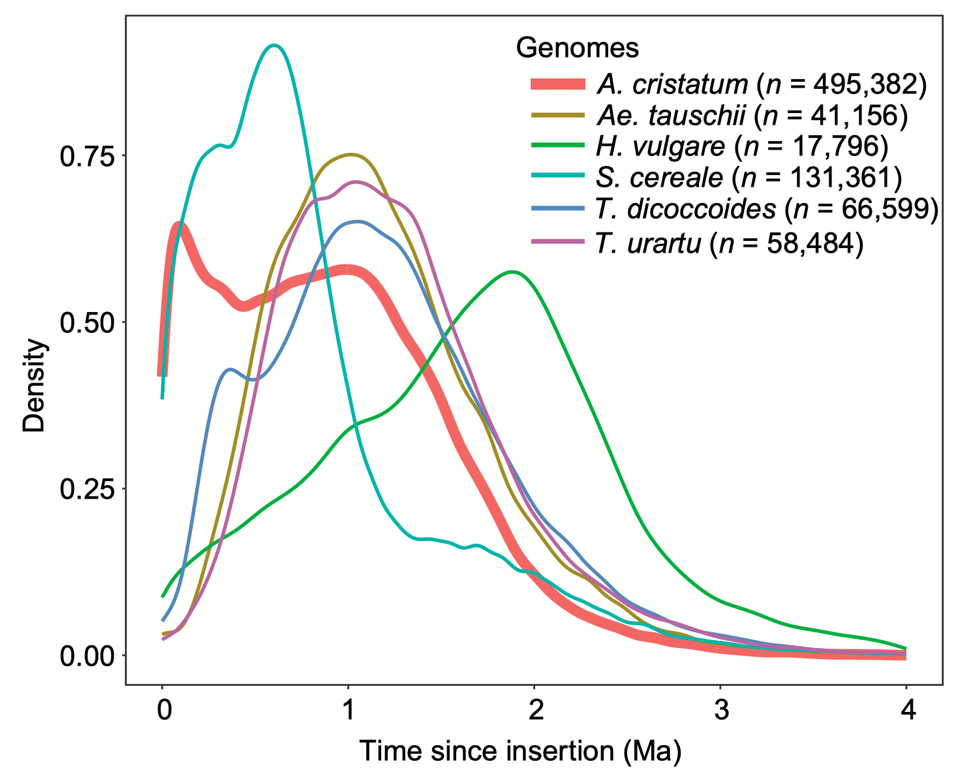

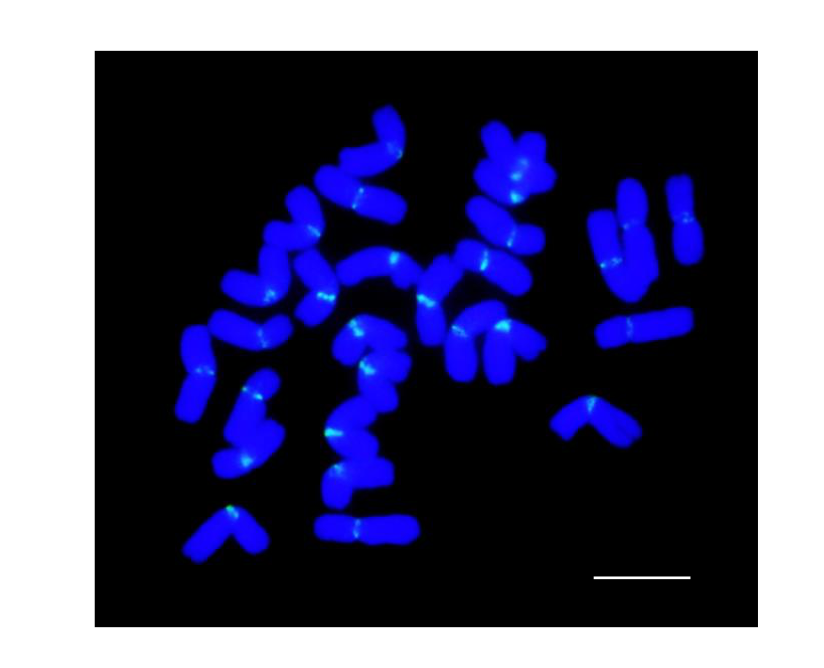
**Supplementary Fig. 4. Temporal patterns of LTR-RT insertion bursts in the A. cristatum genome compared to those in Aegilops tauschii, Hordeum vulgare, Secale cereale, Triticum durum B subgenome, and Triticum urartu genome.** The number of intact LTR-RTs analyzed for each species is indicated in parentheses.

**Supplementary Fig. 5. Chromosomal localization of the partially homologous sequence motif (Oligo-AcCR2) from the *unnamedfam6* retrotransposon using fluorescence in situ hybridization.** Hybridization signals were exclusively detected at the centromeres. Scale bar, 10 μm.

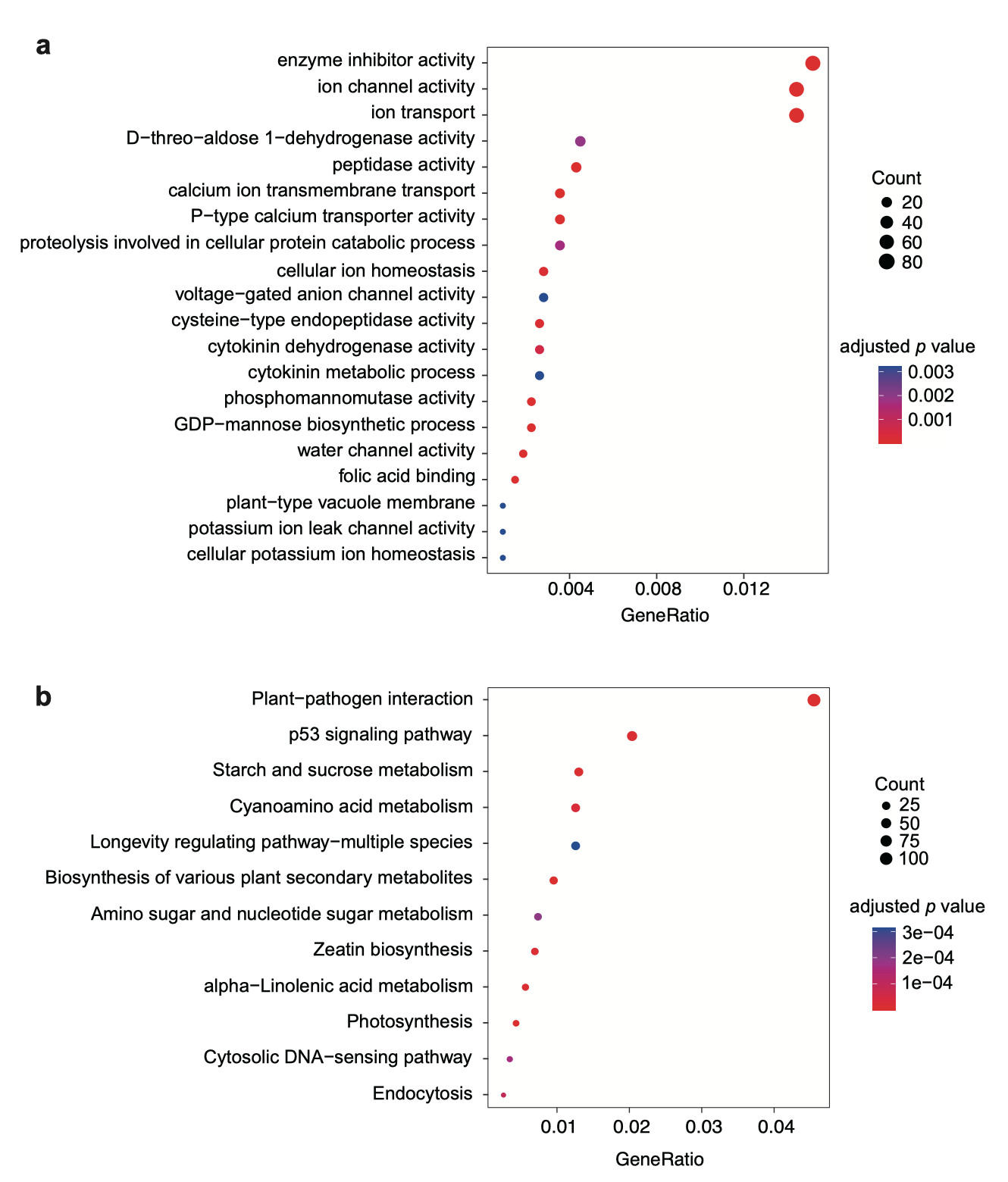
**Supplementary Fig. 6 Enriched GO terms (a) and KEGG terms (b) for *A. cristatum*-specific genes.** *P* values were adjusted using the Benjamini–Hochberg method. GeneRatio represents the proportion of genes associated with a given term relative to the total number of related genes.

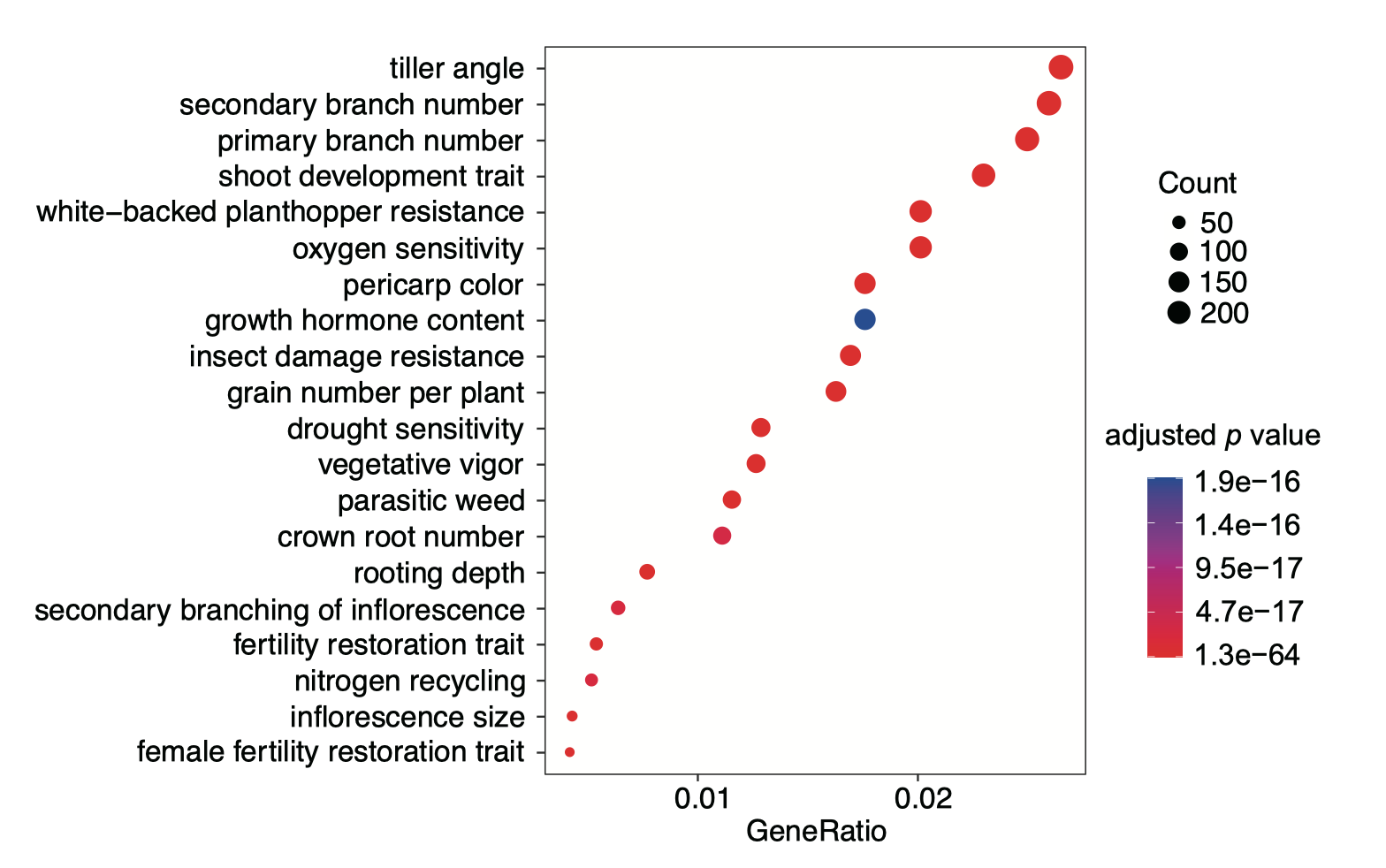
**Supplementary Fig. 7. Enriched TO terms for rapidly expanded genes in *A. cristatum*.** *P* values were adjusted with Benjamini–Hochberg method. GeneRatio represents the proportion of *A. cristatum*-expanded genes associated with a given term relative to the total number of related genes.

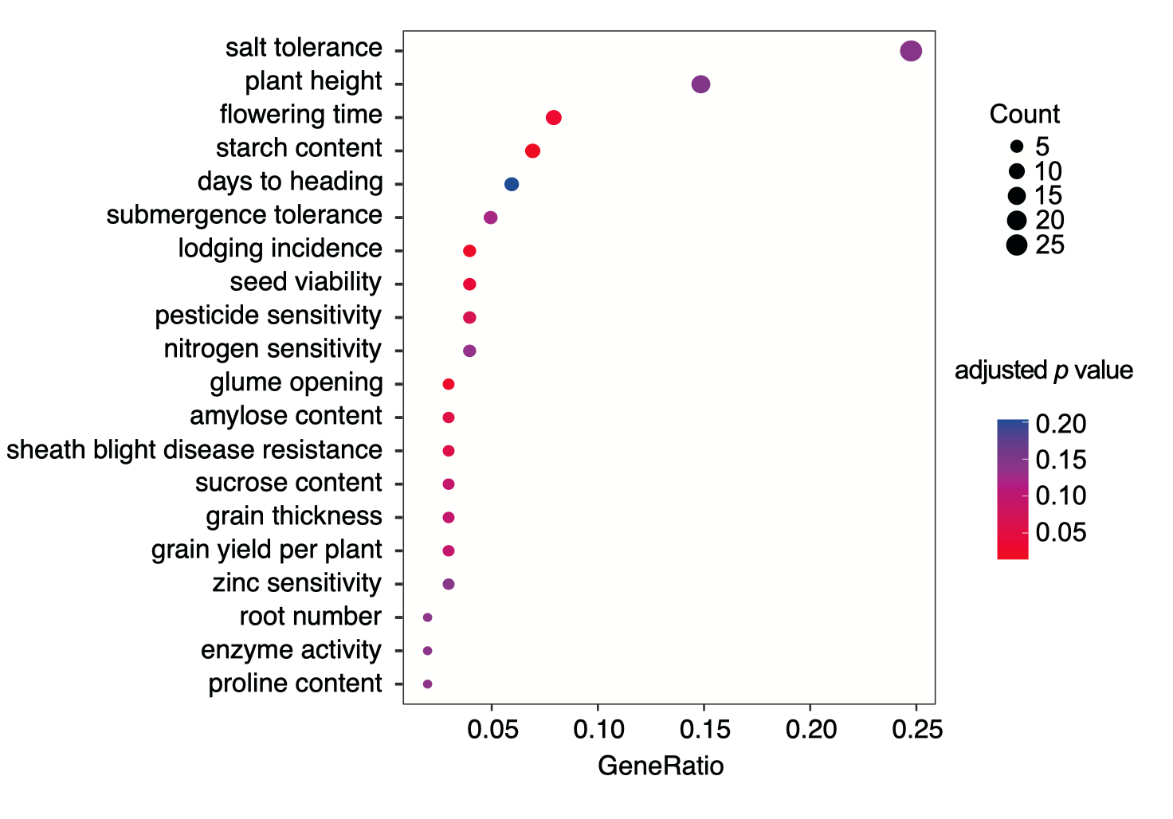

**Supplementary Fig. 8. Enriched TO terms for positively selected genes in A. cristatum.** *P* values were adjusted using the Benjamini–Hochberg method. GeneRatio represents the proportion of positively selected genes in A. cristatum associated with a given term relative to the total number of related genes.

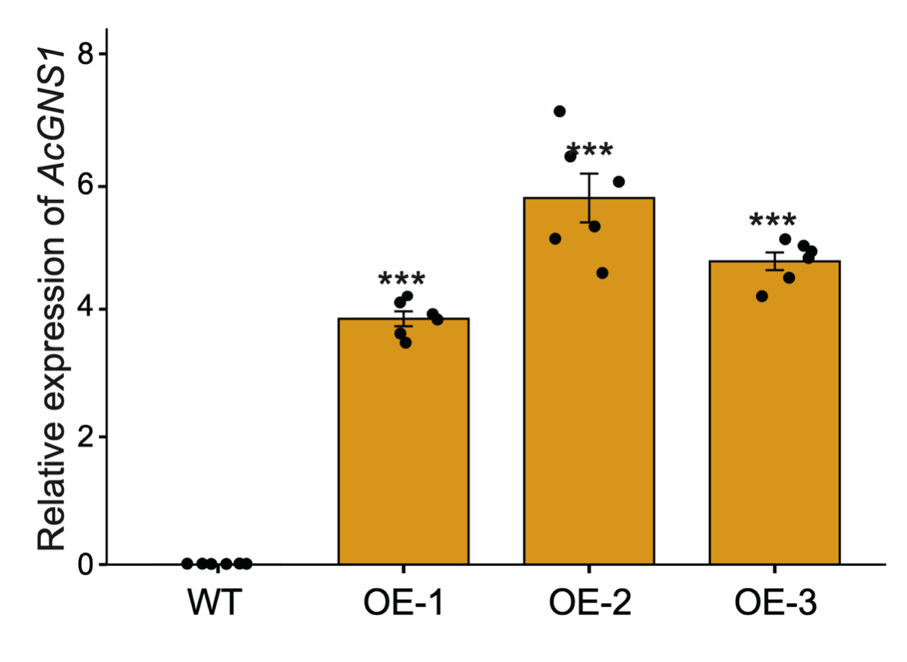
**Supplementary Fig. 9. Comparison of expression levels between wild-type Fielder and three independent *AcGNS1-*overexpressing lines.**

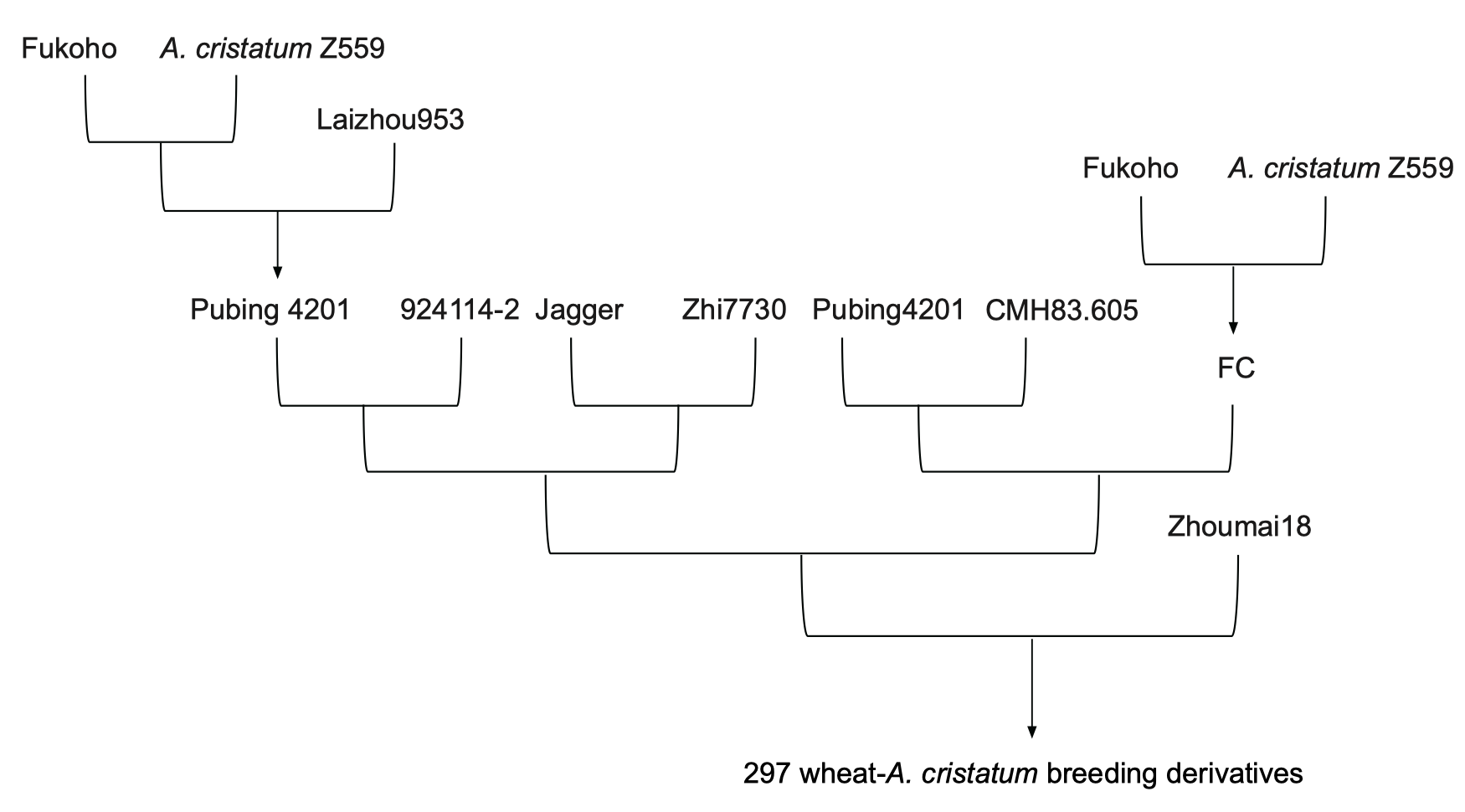

**Supplementary Fig. 10. The pedigree of the 297 wheat-A. cristatum breeding derivatives.**

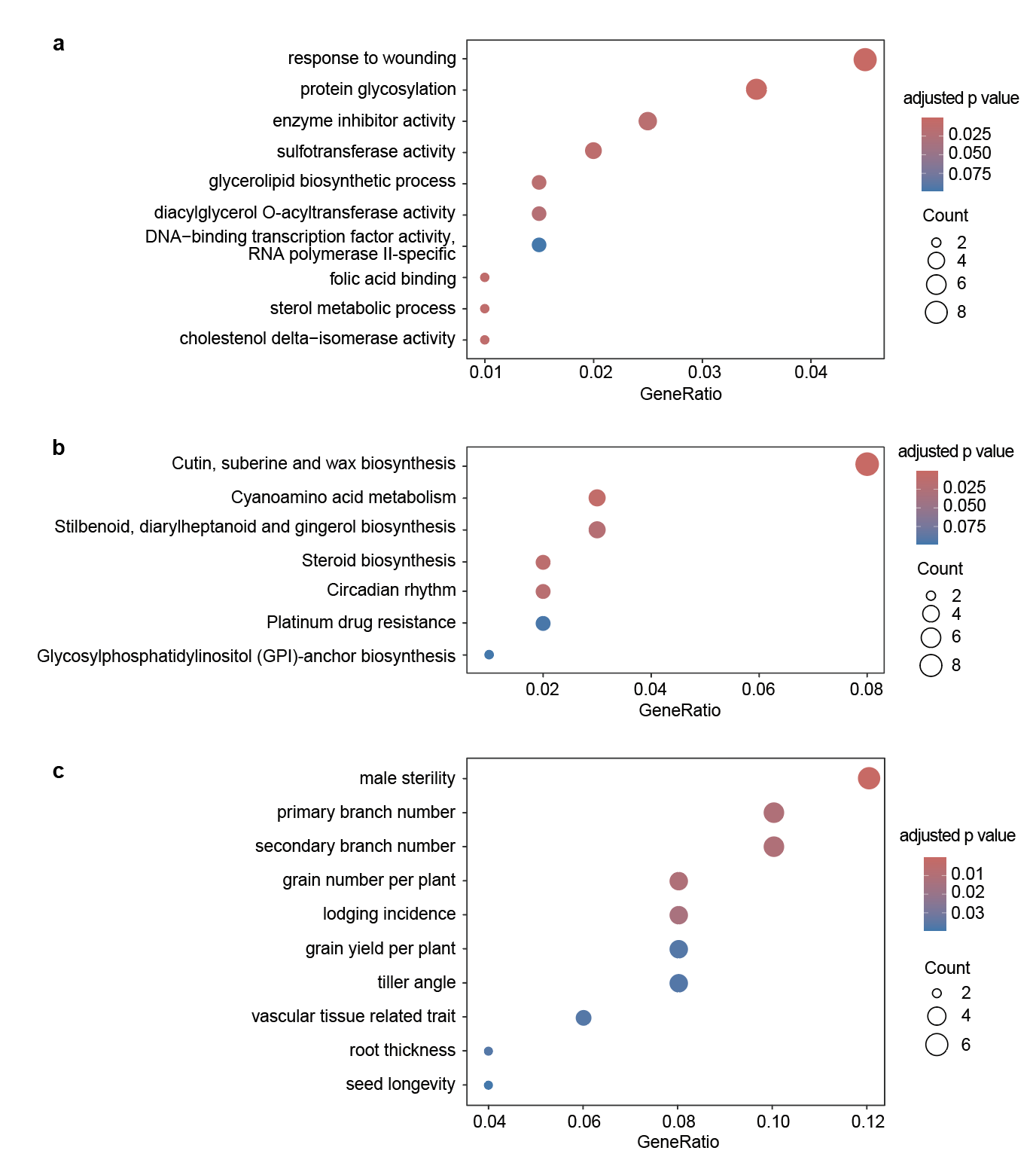
**Supplementary Fig. 11. Enriched GO (a), KEGG (b), and TO (c) terms for A. cristatum alien genes identified in 297 wheat-A. cristatum breeding derivatives.** *P* values were adjusted using the Benjamini–Hochberg method. GeneRatio represents the proportion of genes associated with each term relative to the total number of related genes.

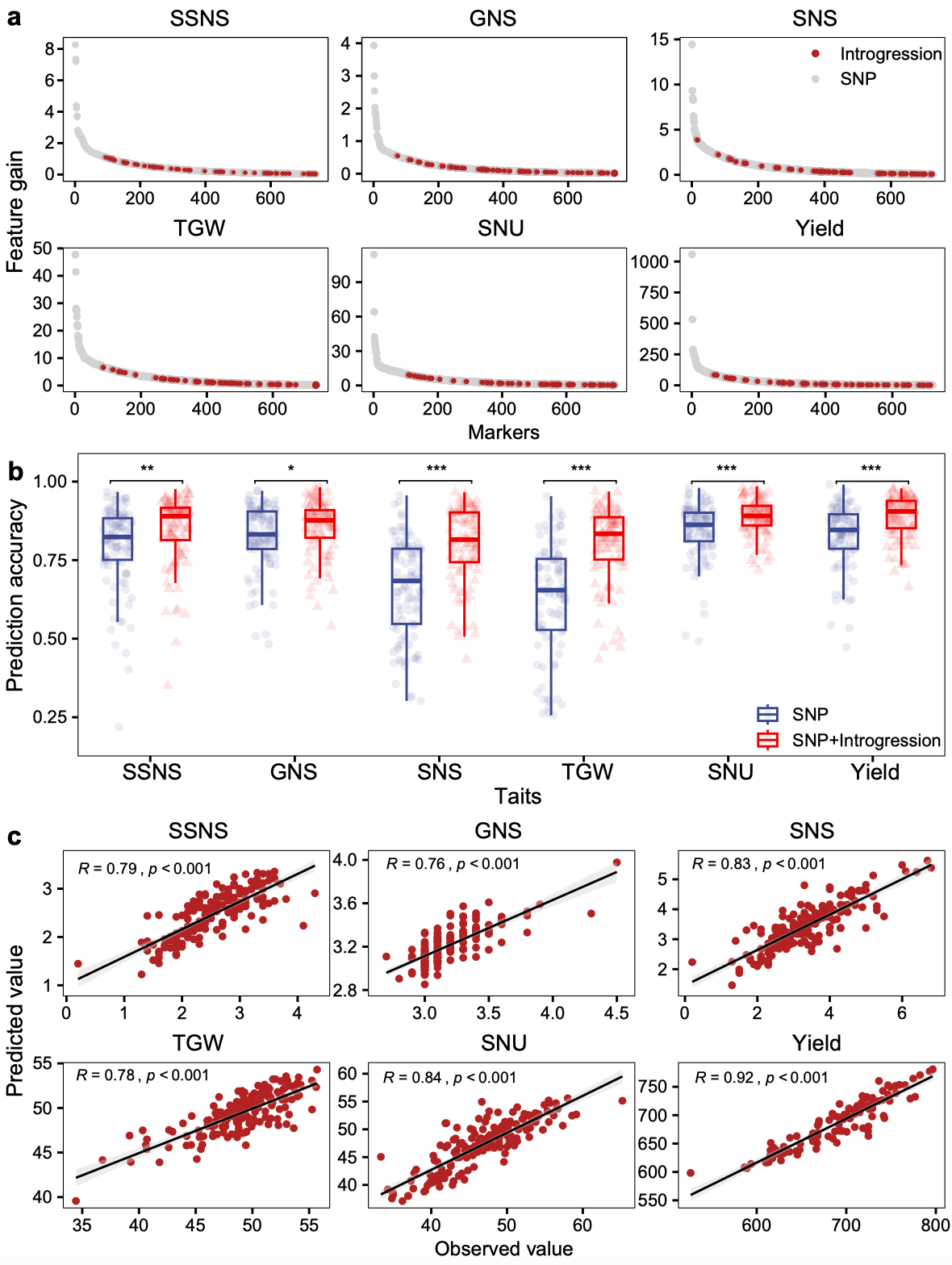
**Supplementary Fig. 12. Development of GS models to predict yield-related traits in wheat-*A. cristatum* breeding derivatives.** **a,** Top features (introgression segments and SNPs) selected that best predict each of the 6 yield-related traits using CropGBM. SNP features are marked in grey while introgression markers are marked in red. **b,** Precision of different phenotypes with different subsets of markers. **c,** Prediction accuracy (measured as Pearson Correlation Coefficient) of trained GS models in hold-out test datasets in each of the six traits.
